# SCALLOPS: a scalable, integrated computational framework for Optical Pooled Screens

**DOI:** 10.64898/2026.05.22.727250

**Authors:** Joshua Gould, Jose Sergio Hleap, Ping Wu, Takamasa Kudo, Jane Guan, Anqi Zhu, Eric Lubeck, Xing-Yue Monica Ge, Aaron Archer Waterman, Tommaso Biancalani, Orit Rozenblatt-Rosen, Ciara Metcalfe, Avtar Singh, David Richmond, Aviv Regev, Bo Li

## Abstract

Optical pooled screens (OPS) link pooled genetic perturbations to high-dimensional image-based phenotypes at scale, but their widespread adoption is hindered by computational bottlenecks in processing terabyte-scale, multimodal image data. We present SCALLOPS, a unified, modular, and cloud-native computational framework that overcomes these bottlenecks. SCALLOPS implements a “well-centric” processing strategy that integrates robust stitching with a non-linear two-stage registration strategy, enabling accurate alignment of multi-magnification images, reliable single-cell genotype–phenotype linkage, and efficient morphological feature extraction. Benchmarking with public and newly-generated datasets demonstrated SCALLOPS’ superior performance over existing solutions. Crucially, SCALLOPS uniquely enables robust processing of 4x magnification *in situ* sequencing data, accelerating image acquisition by around six-fold. We applied SCALLOPS to an optical pooled screen investigating the estrogen receptor (ER) degrader vepdegestrant in a breast cancer cell line, successfully recovering its known mechanism of action, highlighting the value of OPS in translational research. SCALLOPS provides a scalable end-to-end solution, making large-scale OPS routine.

---

Optical pooled screens (OPS) have emerged as a transformative technology that links genetic perturbations with high-dimensional image-based cellular phenotypic readouts at scale. Unlike traditional arrayed approaches^1^ that are highly resource-intensive and have limited scalability, OPS employs targeted *in situ* sequencing (ISS) of expressed perturbation barcodes to connect genotypes and image-based phenotypes from each single cell within a pooled library^2^. OPS has been successfully applied to study a range of biological systems, including perturbation effects on immune signaling^2,3^, antiviral response^4,5^, unfolded protein response^6^, and synaptogenesis^7^. Coupled with Cell Painting^8^ or protein staining^2,4,5,9,10^, OPS also enables probing of perturbation effects on cell morphology^4,11–13^ or multiplex molecular phenotypes^4,5,9,10^. While the original OPS method faced challenges in robustly identifying perturbation barcodes in primary cells and tissues^10,14^, recent methodological advancements^10,14,15^ have addressed these limitations. For example, PerturbView^10^ and NIS-seq^14^ innovatively uncouple perturbation detection from barcode expression level via *in vitro* transcription, enabling flexible and highly multiplexed phenotypic readouts across diverse systems. Together, these innovations enable genome-wide imaging screens to be conducted rapidly and cost-effectively across a broad range of model systems, including cell lines, primary cells, and *in situ* in animal models, more scalably and cost-effectively than Perturb-Seq screens that rely on single cell or spatial profiling.

Despite these advantages, the immense data volume generated by OPS methods, often at terabyte-scale with respect to raw images for genome-wide screens, creates three computational bottlenecks that currently limit their practical application. The first is accurate registration of low-magnification ISS images with their corresponding high-magnification phenotypic images, a non-trivial alignment required to ensure precise genotype-phenotype linkage at the single-cell level. The second is efficient computation and extraction of thousands of complex morphological features from millions of cells for cell-painting-like applications. The third is reliably scaling up image processing for handling terabyte-scale image volumes. Taken together, these have posed a critical barrier to the widespread adoption and routine use of genome-scale OPS.

Current computational solutions for OPS, while functional, generally lack the integration, efficiency, and robustness required for broad adoption. Seminal studies^2,9,16^, and recent large-scale screens^3–5,11,13^, have presented end-to-end workflows, often built on platforms such as Snakemake^17^. However, these implementations are highly customized and built by piecing together a collection of disparate tools: custom scripts for barcode decoding, separate utilities for registration, and imports of functions for feature extraction from general-purpose software like CellProfiler^18^. This disjointed approach is difficult to replicate, extend, and often introduces technical bottlenecks. For instance, cross-magnification registration frequently relies on brittle, non-generalizable scripts to align fiducial ‘anchor points’, which requires manual inspection and can fail easily with modest changes in imaging conditions or biological systems. Moreover, these pipelines are often computationally inefficient, with one study reporting processing times on the order of weeks (≥2) for a single genome-scale dataset^13^. The emerging efforts to collect large scale foundational datasets to build foundation models of cell and tissue biology will require multiple genome-scale (and larger) screens across multiple systems under multiple conditions, necessitating correspondingly efficient and robust computation. Thus, a critical need remains for a unified, efficient, and modular framework that provides robust, scalable solutions for all core OPS analysis steps.

Here, we introduce SCALLOPS (**Scal**able **L**ibrary for **O**ptical **P**ooled **S**creens), a unified, high-performance, and modular computational framework that directly overcomes these critical bottlenecks. To link the phenotype and genotype information for each single cell at different magnifications with high accuracy, SCALLOPS provides fully automated, general-purpose stitching and registration modules and achieves high-fidelity alignment of multi-magnification images at the well level. To enable efficient morphological feature extraction, SCALLOPS employs a parallelized, out-of-core image processing pipeline, with targeted optimizations for the most computationally intensive features. To ensure reliable massive-scale processing of terabytes of image data, we encapsulate SCALLOPS as a cloud-ready solution using the Workflow Description Language (WDL), allowing users to deploy either the full pipeline or customizable WDL tasks efficiently at scale on multiple cloud providers. Critically, unlike previous monolithic or disjointed pipelines, SCALLOPS is fully modular. This design empowers researchers to run the complete end-to-end workflow or to flexibly integrate individual modules, such as the registration or decoding components, into their own custom pipelines.

Through extensive benchmarking on public data, we demonstrate that SCALLOPS is significantly more accurate and computationally more efficient than existing approaches, providing a powerful, accessible tool to make genome-scale optical pooled screens truly routine and robust. Building on the initial validation by Kudo et al.^10^, we demonstrate the first successful multicycle decoding of PerturbView barcodes at 4x magnification, highlighting the potential to accelerate OPS data generation by reducing ISS imaging time around six fold. Finally, by applying SCALLOPS to a multimodal OPS screen that measures several key protein and mRNA phenotypes for estrogen receptor (ER) biology, we validated the known mechanism of action for vepdegestrant, an ER degrader, underscoring the profound potential of OPS in translational research.

## Results

### SCALLOPS is a scalable, versatile end-to-end image processing toolkit for OPS data

To overcome the significant computational bottlenecks associated with analyzing massive, terabyte-scale Optical Pooled Screens (OPS) datasets, we developed SCALLOPS, a scalable, end-to-end and modular toolkit (**Fig. 1a**). SCALLOPS leverages Dask for out-of-core computation and robust parallelization^19^, enabling the reliable processing of large-scale data. Building on this scalable foundation, SCALLOPS introduces three core innovations: (1) efficient stitching to correct microscopic stage drift, (2) robust registration of images acquired at multiple magnifications, and (3) streamlined parallel extraction of cell morphological features. The first two innovations address the registration hurdle and the last one tackles the feature extraction bottleneck.

**Figure 1.**
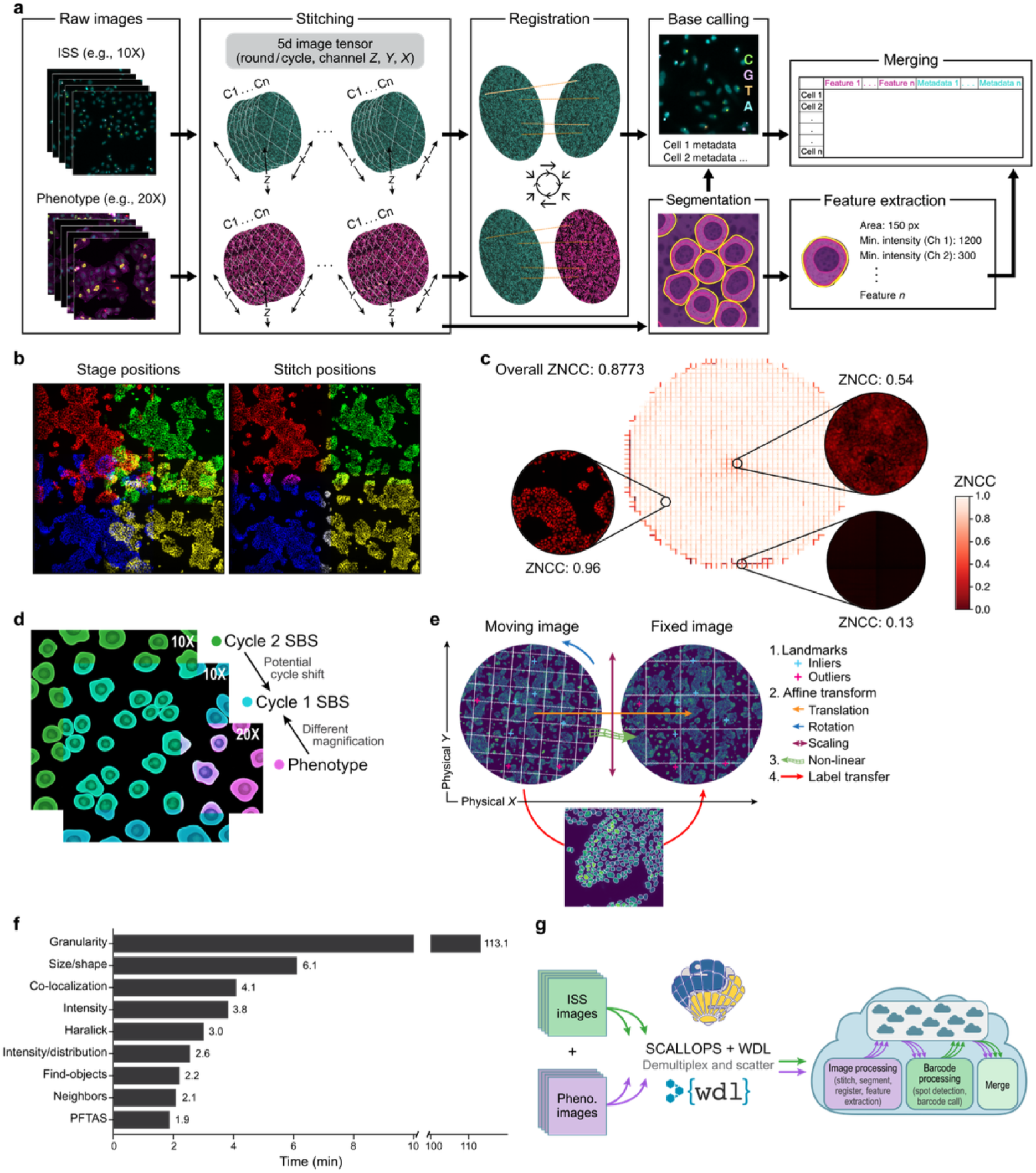
SCALLOPS: a scalable, versatile end-to-end image processing toolkit for OPS data. **a,** SCALLOPS workflow. From left: Raw 2D image tiles from multiple imaging cycles, including both ISS cycles (far left, top) and phenotype channels (far left, bottom) are stitched into a seamless 5D image tensor (Round/cycle, Channel, Z, Y, X) (second left), followed by registration of images from different phenotypic rounds and ISS cycles to correct for spatial shifts (middle). Individual cells are then segmented (second from right, bottom), followed by base calling to determine genotypes (second from right, top), and feature extraction from each cell (bottom right), with all information merged into a single-cell feature matrix for downstream analysis (top right). **b,** Representative example of the stitching module. Image tiles pseudo-colored by original stage positions before alignment (left) and after stitching (right). **c,** Quantitative assessment of stitching accuracy. Center: Zero-normalized cross-correlation (ZNCC) score across the overlapping regions of all adjacent tiles in a well. Insets: stitched images and associated ZNCC in regions with low (bottom right), moderate (left) and very high (top right) cell density. **d,** Illustration of typical alignment challenges between different imaging rounds. Tiles illustrating differences in magnification (10x vs 20x), and physical shifts between an SBS cycle (Cycle 2) and a phenotype stain. Tiles are shown in physical space (μm) and not in image space (pixels). **e,** Multi-stage image registration strategy. The pipeline registers a ‘moving’ image (top left; e.g., phenotype, subsequent SBS cycles, etc.) to a ‘fixed’ reference (top right, usually SBS cycle 1) by applying, in order, landmark detection (1), global affine transformations (2), and non-linear transformation (3). The precise registration then enables the accurate segmentation label transfer (4) from one imaging modality to another. **f,** Efficient feature extraction. Total processing time (x axis) and per-category breakdown (bar labels) for extracting over 1,000 features across multiple feature categories (y axis) from a full well of phenotypic data (20x magnification) of the PERISCOPE A549 screen^13^. **g,** Cloud-native execution workflow. Large-volume input data (left) are demultiplexed and "scattered" across dynamically provisioned Virtual Machines (VMs, right).

### A robust stitching engine assembles overlapping tiles from up to millions of cells

OPS experiments image hundreds to thousands of overlapping tiles to cover a single well containing from several thousands (in high well number plates) to millions of cells (in e.g. 6-well plates). We introduce a robust stitching engine to assemble these tiles into a high-resolution, well-level representation. This well-centric strategy prevents cell double-counting in overlapping regions and obviates the need for tile-to-tile correspondence required by previous pipelines^16,20^. Consequently, this relaxes constraints on microscope selection by removing the need for high-precision stages or manual alignment between acquisition cycles, while providing the global context necessary to accurately register images across different magnifications.

Specifically, we developed a new SCALLOPS stitcher that builds upon the adjacency graph framework proposed by FijiIS^21^, MIST^22^ and ASHLAR^23^, by taking the imperfect stage positions as priors and outputting refined coordinates that align most overlapping regions accurately (**Fig. 1b**). To this end, the SCALLOPS stitcher first maps stage positions (in microns) to its internal pixel-based coordinate system, and optionally corrects each tile for radial distortion^24^ (default: on). It then evaluates tile pairs to construct an adjacency graph based on stage-position-guided overlaps, with edge weights defined by zero-normalized cross-correlation (ZNCC) scores (**Methods**). We chose to evaluate overlapping regions with ZNCC rather than normalized cross correlation (NCC), because it was less sensitive to the mean intensity differences between adjacent tiles (**Extended Data Fig. 1a**). SCALLOPS estimates a null distribution of ZNCC score from randomly sampled non-overlapping tile pairs, and filters the edges based on the null distribution and a user-defined shift threshold. It then identifies connected components, and determines optimal tile positions within each component, using a global optimization algorithm initialized with a spanning-tree-based greedy solution. Finally, it projects all components into a common coordinate system using a linear regression model trained on the largest component (**Extended Data Fig. 1b**, **Methods**). Once the stitch positions are finalized, the SCALLOPS stitcher fuses individual tiles into a well-level mosaic image (**Methods**), which is used for downstream analysis. We implemented parallel writing of mosaic images, which significantly accelerates the runtime of our stitcher, especially for datasets with multiple channels (**Extended Data Fig. 1c**).

The SCALLOPS stitcher introduces three innovations over previous works^21–23^. First, it uses both cross correlation^25^ and phase correlation^26^ to determine the best translation between adjacent tiles (**Methods**), based on our observation that each method performs best for a subset of cases, but neither is consistently the best (**Extended Data Fig. 1a**). Second, it uses the weighted *L*2 norm as the objective function for global optimization (**Methods**), where the translation of each overlapping tile pair is weighted by its ZNCC score. Using a weighted *L*2 norm allows us to emphasize high-quality overlapping pairs while de-emphasizing pairs that share few cells and often exhibit low ZNCC scores. Lastly, it is able to automatically determine and correct radial lens distortion (barrel or pincushion) at the tile level before stitching (**Methods**).

Benchmarked against the MIST ground truth dataset, SCALLOPS (without radial correction) outperformed MIST and ASHLAR in the *D*_*err*_ metric, while achieving comparable performance across the remaining two metrics **(Extended Data Fig. 1d, Methods).** Further evaluation on four wells of OPS data using the overall ZNCC score showed that SCALLOPS with radial correction had the highest fidelity, followed by the uncorrected version, both of which surpassed ASHLAR **(Extended Data Fig. 1e)**.

Another unique feature of the SCALLOPS stitcher is the diagnostic visualization for stitching quality (**Fig. 1c**), which highlights how well each overlapping region is aligned using the ZNCC scores. We additionally summarize ZNCC scores of all overlaps into a single, well-level overall ZNCC score, which can be used to compare stitching quality of different stitchers or stitching parameters (**Methods**). SCALLOPS’ diagnostic visualization helps users identify potential data quality issues, such as severely out-of-focus images, empty or missing tiles, and stage position mismatches, as high-quality tile pairs typically yield high ZNCC scores (**Fig. 1c**).

### A two-stage strategy registers images across cycles, modalities, and magnifications

SCALLOPS employs a two-stage approach to register images across different ISS cycles, modalities, and magnifications, which often exhibit significant rotation and physical shifts (**Fig. 1d, Extended Data Fig. 2a,b**). The registration module first corrects for large-scale misalignments through a landmark-based initialization, identifying corresponding points via template matching^27^ and RANSAC filtering^28^ to compute a coarse transformation. Next, it performs global affine transformations to resolve linear shifts (translation, rotation, scaling and shearing) and a final non-linear transformation to correct local distortions using the Insight Segmentation and Registration Toolkit (ITK) via the elastix library^29^ (**Fig. 1e**). SCALLOPS provides predefined parameter settings for rigid, affine, similarity, and non-linear B-spline transformations adapted from common practices^30^, while also supporting fully custom elastix parameters. The final, high-precision transformation can be applied to any image channel or segmentation mask, enabling the accurate transfer of labels (e.g., from a high-quality DAPI mask) to all other cycles for base calling and feature extraction (**Fig. 1e**).

We compared our full registration with translation-only registration on real data (**Methods**) since several tools^13,23,31^ only consider translation for image registration. Our results clearly demonstrated the necessity of applying both linear and non-linear transformations for image registration (**Extended Data Fig. 2d**). A unique feature of SCALLOPS is its ability to assess registration quality at the single-cell level by correlating DAPI intensities from the reference phenotypic and ISS datasets within each cell’s segmented bounding box (**Methods, Extended Data Fig. 2c**). With this feature, users can easily filter out cells with low registration quality, thereby improving overall data quality for downstream analyses.

### Efficient feature extraction and end-to-end OPS workflow with SCALLOPS

After segmentation masks are accurately transferred via registration, SCALLOPS performs efficient feature extraction by leveraging parallelized out-of-core computation and speed-optimized reimplementation of subsets of features. To overcome the memory and CPU-bound limitations of traditional methods, SCALLOPS employs a custom, highly scalable engine built on Dask^19^, with an innovative two-pass approach: it first globally identifies all unique object bounding boxes, then processes the image in parallel, out-of-core blocks, calculating features only for objects intersecting each block. This engine supports all single-mask and neighbor features provided by cp_measure^32^, covering most CellProfiler features across seven categories: SizeShape, Colocalization, Granularity, Intensity, IntensityDistribution, Neighbors, and Texture (**Supplementary Table 1**). In addition, the module also integrates Parameter-Free Threshold Adjacency Statistics (PFTAS)^33^, a set of texture features not covered by cp_measure or CellProfiler. We benchmarked the speed of each SCALLOPS feature category using one well of data from the PERISCOPE^13^ A549 screen (**Methods**) to illustrate its performance (**Fig. 1f**). While SCALLOPS leverages validated algorithms from cp_measure, it has additional performance gains thanks to its custom, optimized implementations for intensity and colocalization features, achieving 173- and 5-fold speedups vs. cp_measure (v0.1.12) for these two categories, respectively (**Extended Data Fig. 2e**).

Beyond these core innovations, SCALLOPS provides several other critical, scalable modules to complete the OPS workflow. For cell identification, it includes robust nucleus and cell segmentation algorithms and leverages Dask to operate entirely out-of-core, efficiently managing memory while processing massive images. This module is also flexible, supporting the use of multiple channels to generate a more accurate composite cell mask. For genotype decoding, SCALLOPS implements an automated workflow that replaces manual peak thresholding by algorithmically determining an optimal peak intensity cutoff based on read quality scores (**Methods**). Finally, it integrates the U-FISH^34^ deep learning model for robust FISH spot detection, enabling accurate RNA expression quantification in different cellular compartments.

To orchestrate end-to-end analysis on the cloud and address massive data volumes, we provide a Workflow Description Language (WDL) workflow. This workflow automates the entire OPS pipeline, orchestrating all SCALLOPS modules—from stitching, registration and segmentation through feature extraction, base calling, and data merging. It is designed for massive parallelization, "scattering" computation across experimental units such as plates or wells (**Fig. 1g**). This design enables seamless, on-demand resource provisioning in the cloud, as each task specifies its precise computational requirements (e.g., CPU, memory, and disk), such that cloud execution engines can automatically allocate the necessary resources, resulting in a reproducible, automated, and highly scalable solution for processing terabyte-scale OPS datasets. Using these cloud-based workflows, we processed end-to-end in only half a day all nine plates of PERISCOPE A549 screen data into analysis-ready feature vectors from over 12.8 million cells, compared to at least 2 weeks by the PERISCOPE pipeline^13^ (**Extended Data Fig. 2f**).

Together, this combination of a scalable architecture, efficient out-of-core processing, and a modular, cloud-ready workflow positions SCALLOPS as a robust solution, enabling researchers to overcome the primary computational and analytical bottlenecks in processing diverse, large-scale OPS datasets from raw images to final feature tables.

### End-to-end SCALLOPS pipeline for standard OPS and PerturbView

To demonstrate SCALLOPS’ performance on standard OPS data, we applied it to reprocess all 9 plates of data from the PERISCOPE A549 screen^13^. We compared SCALLOPS to the original analysis based on correlation with next-generation sequencing (NGS) counts extracted from genomic DNA and the number of usable cells at the well level (**Fig. 2a**). For SCALLOPS, we first applied a novel single-cell level filter based on overlaps with the stitching boundary or the registration quality between phenotype and ISS (**Methods**), and then tested three *post hoc* barcode purity filtering rules with increasing stringency, all restricted to barcodes matching the whitelist (**Methods**): max count rule, whereby each cell is assigned to the barcode of most counts; 50% rule, where we only keep cells where the most abundant barcode is >50% of total barcode counts; and 100% rule, where we only keep cells where all barcodes are identical. In all cases, we filtered out cells without any barcode assigned. The original analysis applied an approach similar to the 100% rule.

**Figure 2.**
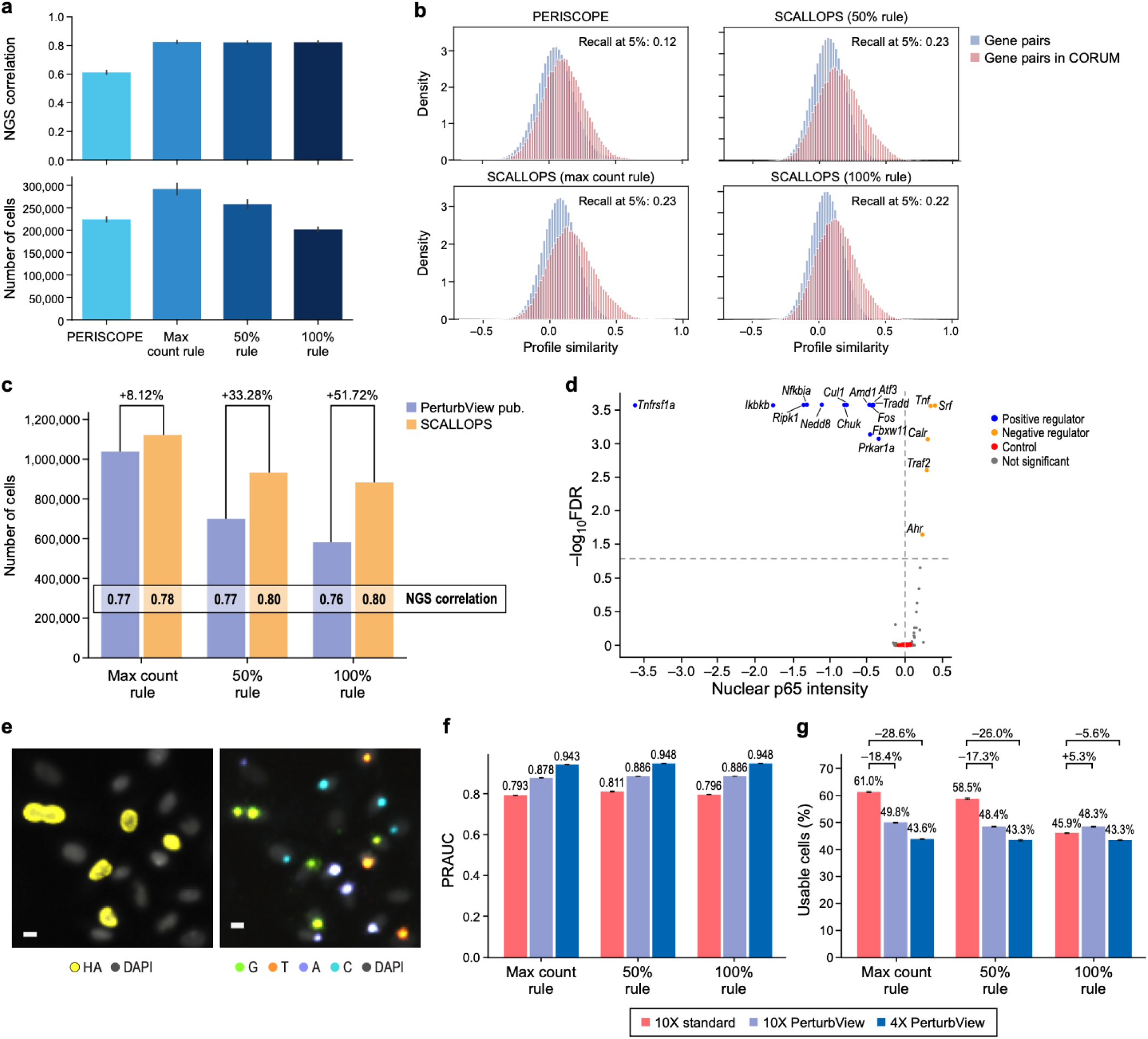
Performance evaluation of SCALLOPS across OPS and PerturbView datasets and a frameshift reporter “ground truth” screen. **a,b,** SCALLOPS performs well on PERISCOPE’s A549 OPS data. **a,** Correlation with NGS counts (top, y axis) and number of recovered cells (bottom, y axis) using the original analysis method (PERISCOPE) or SCALLOPS with max count, 50%, and 100% rules (x axis) at the well level. Error bars: standard error of the mean (SEM) across wells. **b**, Distributions of cosine similarity (x axis) between the phenotypic profiles of random gene pairs (blue) or gene pairs in CORUM 4.0 protein complexes (red) for the original PERISCOPE analysis or for SCALLOPS with either of three filtering rules. To right: Recall at 5% value (**Methods**). **c,d,** SCALLOPS performs well on an NF-*κ*B PerturbView screen in primary mouse bone marrow-derived macrophages (BMDMs). **c**, Number of usable cells (y axis) recovered by the original analysis^10^ or by SCALLOPS with either of three filtering rules (x axis). Correlations with NGS counts from the plasmid libraries are shown within each bar. **d**, Significance (-log_10_(FDR), y axis) and effect sizes (defined as the median of guide-level medians of normalized nuclear p65 intensity, **Methods**, x axis) for each gene (dot) under TNF-*α* stimulation, colored as positive regulators (blue; FDR<0.05), negative regulators (orange; FDR<0.05), non-significant (grey; FDR≥0.05) or nonessential gene controls (red). **e-g,** SCALLOPS’ performance on frameshift reporter (“ground truth”) screen. **e,** Example images of the same cells (nuclei stained by DAPI, grey) with HA epitope stain (left, yellow) and with cycle 1 ISS stain (right, G,T,A,C). All targeting guides start with G (green) and all nontargeting guides start with T (orange), A (purple) and C (blue); scale bar, 10 μm. **f,g,** Cell-level barcode calling accuracy (PRAUC, y axis, **f**) and percent of usable cells (y axis, **g**) at the well level for SCALLOPS’ analysis of 10x Standard OPS (red), 10x PerturbView (purple), and 4x PerturbView (blue) (x axis) using each of the three rules (x axis). Error bars: SEM across wells.

SCALLOPS had much higher correlations with NGS counts than the original analysis comparably across all filtering rules (**Fig. 2a**, top), recovering more cells than the original analysis with the max count and 50% rule, and somewhat fewer with the 100% rule (**Fig. 2a**, bottom). All three SCALLOPS analyses also exceeded the original one based on their recall of perturbed gene pairs with similar perturbation effects that are members in the same protein complex in the CORUM database^35^ (defined as the fraction of top 5% gene pairs ranked by cosine similarity that are in CORUM among all gene pairs both in the screen and CORUM^36^; **Fig. 2b**, **Methods**). This suggests that SCALLOPS produced better quality cells than the PERISCOPE pipeline and that the 100% rule might be too stringent for this screen.

We next evaluated SCALLOPS’ performance on a PerturbView screen of mouse primary bone marrow-derived macrophages (BMDM)^10^ perturbed for components on the NF-*κ*B pathway and stimulated with LPS, TNF-*α*, or IL-1*β*. Applying the same barcode purity rules to both SCALLOPS and the PerturbView pipeline, SCALLOPS preserved more cells under all three filtering criteria with better correlations with NGS counts from the plasmid library (**Fig. 2c**). As NGS count correlation improves between the 50% rule and max count rule, but not further by the 100% rule, the 50% rule may be the optimal balance for *post hoc* cell filtering. Leveraging SCALLOPS’ single-cell level filters led to a further, albeit modest, improvement in the correlations with NGS counts compared to the already enhanced SCALLOPS baseline, with an expected reduction in the number of usable cells (**Extended Data Fig. 3a-c**).

The 54 hits obtained using SCALLOPS-based analysis (50% rule and single-cell level filters, **Methods**) included 53 of 64 hits identified in the original analysis^10^, including established NF-*κ*B pathway regulators^37^ such as *Tnfrsf1a*, *Ikbkb*, *Ripk1*, *Nfkbia*, *Nedd8*, *Chuk*, *Tradd*, *Traf2*, *Il1r1*, *Myd88*, *Irak4*, *Irak1* and *Traf6*, indicating high reproducibility (**Fig. 2d**, **Extended Data Fig. 3d,e**, **Supplementary Table 2**). Importantly, 8 of the 11 hits unique to the original analysis were weak (adjusted *P* ≥ 0.01), implying that their exclusion reflects the more rigorous data processing inherent to SCALLOPS. We further investigated *Map3k7* under LPS stimulation, the strongest hit unique to the original analysis. In the original study, one of the four targeting guides (blue) displayed a divergent cumulative distribution function (CDF) profile (Extended Data Fig. 3d from Kudo et al.^10^), indicative of a potential outlier. However, following SCALLOPS processing, the CDF profile of this guide aligned with the other three guides (**Extended Data Fig. 3f**), demonstrating that SCALLOPS effectively filters out aberrant data points. Additionally, the original PerturbView study highlighted *Prkar1a* as a negative regulator under IL-1*β* stimulation; we reproduced this and additionally found that *Prkar1a* was a positive regulator under TNF-*α* stimulation (**Fig. 2d**, **Extended Data Fig. 3g**). A previous *PRKAR1A* depletion study in the human H295R cell line suggested *PRKAR1A* knockdown promoted the nuclear translocation of p50 protein, but not p65, in unstimulated cells^38^. Taken together, our results demonstrate that SCALLOPS is a reliable tool for the end-to-end processing of both standard OPS and PerturbView data.

### Efficient decoding of PerturbView barcodes at 4x magnification through a frameshift reporter screen

To evaluate the quality of guide assignment directly, we performed a dedicated frameshift reporter screen using the A549 cell line^10^ (**Fig. 2e**), with a library of 5 targeting and 5 nontargeting guides. Upon activation, each targeting guide can trigger a frameshift and thus turn on the expression of a hemagglutinin (HA) epitope^2^. The HA epitope serves as a proxy of the ground-truth measurement of the perturbation outcome (**Fig. 2e**, left), which can in turn be compared with the cell-level guide assignment inferred from ISS data (**Fig. 2e**, right). Our screen encompassed three settings: standard OPS with ISS imaged at 10x magnification, and a PerturbView with ISS where the same cells were imaged at 10x and 4x magnifications, each performed in triplicate. Unlike previous frameshift reporter assays^2,10^, we imaged HA epitopes (phenotypic data) at 20x magnification and conducted 8 rounds of ISS for barcode calling to ensure that the screen closely matches real OPS screens, with the need to register the same cell imaged at different magnification. These data are thus a foundational benchmark for future OPS computational tools.

We benchmarked SCALLOPS on the frameshift screen by applying the single-cell level filters followed by one of the three barcode purity rules. We used a well-level Area Under the Precision–Recall Curve (PRAUC)^39^ metric to compare the guide assignment with the HA epitope, thus avoiding the need to determine a cutoff for HA epitope activation^10^. We defined precision as the fraction of cells with both targeting guides and HA epitopes (true positives) among all cells with targeting guides and recall as the fraction of true positives among all cells with any guides and HA epitopes. Thus, our assessment focused on the quality of barcodes called among all cells with detected barcodes (a distinct assessment than that in Kudo et al.^10^).

SCALLOPS performed well across all settings with the 50% rule achieving the best PRAUC value within each setting (**Fig. 2f**), consistent with our NGS counts correlation (**Fig. 2a,c)**. While PerturbView enabled more reliable guide assignment than standard OPS, the differences were modest in this assay, likely reflecting the relatively favorable experimental context—a low-plex measurement in a cancer cell line—in which standard OPS is already highly sensitive, whereas PerturbView’s sensitivity advantages are typically more pronounced in challenging cellular contexts and integrations with other assays and stains^10^. Surprisingly, 4x PerturbView had the highest PRAUC value across all settings, likely because spurious spots are invisible at low magnification. Although 4x PerturbView could significantly increase data capture rate for large screens^10^, its low resolution ISS images pose challenges for almost all data processing steps. While ASHLAR, a state-of-the-art image stitcher, failed to stitch the 4x ISS images (**Methods**), SCALLOPS successfully analyzed these data for the first time in a realistic setting where phenotype and ISS are captured at different magnification.

We also assessed the well-level percentage of usable cells, defined as the fraction of cells retained after barcode purity filtration out of the total number of segmented cells after the single-cell level filters (**Methods**). Here, standard OPS retained a higher percentage of usable cells under all filtering rules (**Fig. 2g**), although the three experimental settings yielded near-identical cell fractions with the stricter 50% and 100% rules (**Fig. 2g**). Overall, there is a trade-off between guide assignment reliability and percentage of usable cells across the three settings, such that 4x PerturbView produces the highest quality usable cells, but a reduced yield (**Fig. 2f,g**). Because ISS imaging time governs data capture rate and 4x PerturbView reduces it by around six-fold^10^, it has the potential to significantly boost the usable cell yields for large-scale OPS screens.

### SCALLOPS validates an ER degrader’s mechanism of action through a multimodal optical pooled screen

We applied SCALLOPS to a multiplexed optical pooled CRISPR knockout screen in the MCF7 breast cancer cell line to illustrate the known mechanism of action (MoA) of vepdegestrant^40^, an investigational molecule currently under development for the treatment of ER+ breast cancer. Vepdegestrant is a heterobifunctional degrader that recruits the CRL4-CRBN E3 ligase to ER, leading to its polyubiquitination and proteasomal degradation, a canonical PROTAC mechanism^41,42^ (**Fig. 3a**). The screen library contains 4 sgRNAs for each of 146 genes, manually curated to focus on the ubiquitin-proteasome system (UPS), ER signaling pathway and common tumor suppressors (**Methods**, **Supplementary Table 3**). We performed the screen using an established multimodal optical screening workflow^2,10^ with some modifications (**Methods**). The screen quantified, in the same cells, the ER protein (by immunofluorescence) as well as the RNA transcripts (by FISH) for two ER transcriptional targets, *GREB1* and *CCND1,* and for the ER gene, *ESR1* (**Fig. 3b**).

**Figure 3.**
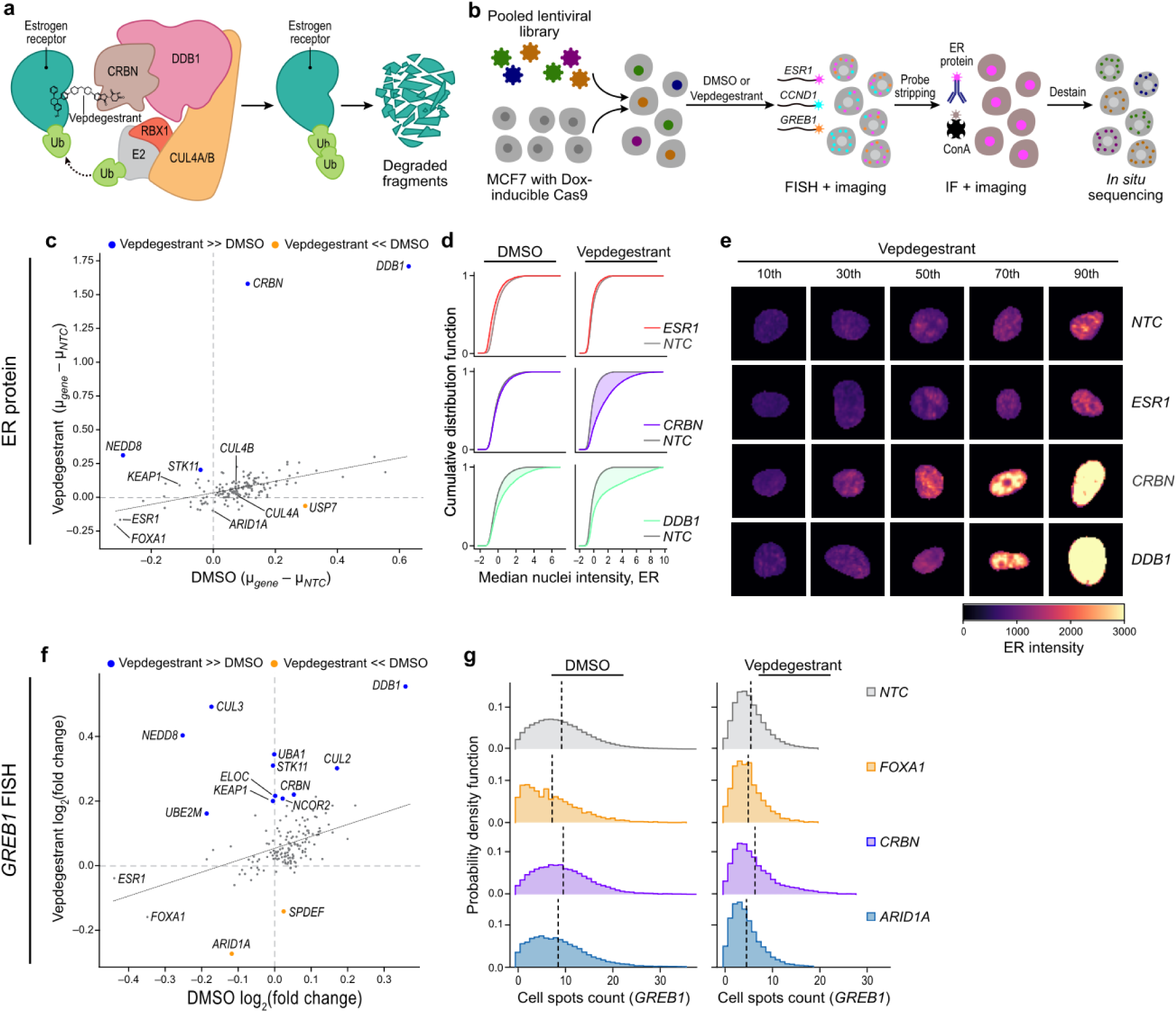
SCALLOPS confirms vepdegestrant’s MoA via a multimodal optical pooled screen. **a,** Mechanism of action for vepdegestrant, a PROTAC degrader. Vepdegestrant recruits the CRL4-CRBN E3 ubiquitin ligase complex (containing *CRBN*, *DDB1*, *CUL4A/B*, and *RBX1*) to the Estrogen Receptor (ER) protein, leading to its polyubiquitination and subsequent proteasomal degradation. **b,** MCF7 multimodal OPS. From left: MCF7 cells harboring doxycycline-inducible Cas9 (bottom left) were transduced with a pooled lentiviral sgRNA library (top left), followed by puromycin selection for 3 days. Cells were then split and treated with DMSO (vehicle) or vepdegestrant (arrow) for 24 hours, and then fixed and profiled sequentially by multiplexed HCR FISH to quantify *ESR1*, *CCND1*, and *GREB1* mRNA transcripts (second from left), immunofluorescence (IF) to measure ER protein levels and ConA for cellular mask (second from right), and finally, ISS to identify the sgRNA in each cell (right). **c,** Treatment-dependent gene-level knockout effects on ER protein abundance. Genes (dots) with significant (FDR≤0.05) effects on ER protein abundance in DMSO (x axis) or vepdegestrant (y axis) treatment. Colored points: hits with higher effect in vepdegestrant (blue) or DMSO (orange) conditions (outside the RANSAC regression 99% prediction interval, dotted black line, **Methods**). Direct regulators of basal ER levels (*ESR1*, *FOXA1*) and other genes of interest (*ARID1A*, *CUL4A/B*, *KEAP1*) are marked. **d,e,** Impact of key regulators. Cumulative distribution functions of ER protein nuclear intensity (**d**) and representative images (**e**, sampled at different percentiles of normalized ER median intensity (local z score), columns) of cells with non-targeting controls (NTC, grey) or with guides targeting *ESR1* (red, top), *CRBN* (purple, middle) or *DDB1* (green, bottom) in DMSO (**d**, left) and vepdegestrant (**d**, right and **e**) treatment. **f,g,** Effects on *GREB1* expression (an ER transcriptional target). **f,** Genes (dots) with significant effects (FDR≤0.05) on *GREB1* expression level in DMSO (x axis) or vepdegestrant (y axis) treatment. Colored points: hits with higher effect in vepdegestrant (blue) or DMSO (orange) conditions (outside the RANSAC regression 99% prediction interval, dotted black line). Direct regulators of baseline ER signaling (*ESR1*, *FOXA1*) are annotated for reference. **g,** Distributions of *GREB1* FISH spot counts per cell (x axis) in cells with NTC guides or with guides targeting specific genes under DMSO (left) or vepdegestrant (right) treatment. Vertical dashed lines: distribution mean.

SCALLOPS processed data was highly reproducible and confirmed the known MoA of the drug. SCALLOPS recovered highly reproducible gene-level effects in biological replicates for both treatment and vehicle conditions (e.g., Pearson’s *r* ≥ 0.97, 0.86, 0.91, 0.94 for the ER protein and *ESR1*, *CCND1*, *GREB1* transcripts respectively, **Extended Data Fig. 4a,b**). Furthermore, knockouts of direct regulators of ER expression, like *FOXA1* and *ESR1*, reduced ER protein levels in both DMSO and vepdegestrant treatment, confirming known biology (**Fig. 3c-e**). Analysis of the knockout gene effects on the ER protein phenotype confirmed the known MoA of vepdegestrant. Specifically, knockout of the core CRL4-CRBN E3 ligase components *CRBN* and *DDB1* prevented vepdegestrant-mediated ER degradation^40,43^ (**Fig. 3c-e**, **Extended Data Fig. 4c,d**). These knockouts did not show vepdegestrant-specific effects on *ESR1* mRNA levels, confirming the drug’s post-translational action (**Extended Data Fig. 4e-g**). Knocking out the cullin scaffolds *CUL4A* and *CUL4B* had much weaker effects, likely reflecting their partial redundancy in MCF7 cells^44^. *CRBN* and *DDB1* knockout also partially rescued *GREB1* expression, confirming that blocking the E3 ligase axis restores some ER-dependent transcription (**Fig. 3f,g**, **Extended Data Fig. 5a,b**). Beyond the core degrader axis, the screen revealed orthogonal regulatory mechanisms. Loss of the SWI/SNF subunit *ARID1A*, which has been linked to endocrine therapy resistance^45^, significantly reduced *GREB1* transcription (vs non-targeting control, NTC) – particularly under the treatment condition (**Fig. 3f,g**) – perhaps consistent with its role in maintaining luminal lineage identity and thus ER competence, when under pressure of endocrine therapy^46,47^.

Our analysis also generated novel hypotheses by implicating the tumor suppressor *STK11* and the ubiquitin-like protein modifier *NEDD8* in ER biology. Both genes exhibited significant treatment-specific knockout effects, with increased ER protein abundance (**Fig. 3c**) and elevated *GREB1* and *CCND1* transcript levels (**Fig. 3f**, **Extended Data Fig. 5**). Additionally, knockout of *KEAP1*, a key negative regulator of the NRF2 pathway, resulted in a treatment-specific increase of *GREB1* transcript levels, accompanied by a modest trend toward increased ER protein abundance and *CCND1* transcript levels (**Fig. 3c,f**, **Extended Data Fig. 5a,b,e**). These results suggest potential roles for *STK11*, *NEDD8*, and *KEAP1* in ER signaling and endocrine therapy resistance. Notably, both *STK11* and *KEAP1* have been previously implicated in endocrine resistance in a functional genomics screen relying on a cell viability readout, but without a direct connection to ER biology^48^. While the new hypotheses uncovered by this screen require extensive follow-up, this work provides a multi-layered portrait of vepdegestrant MoA: it confirms the canonical post-translational degradation, links it to downstream transcriptional consequences, and uncovers orthogonal pathways that may modulate drug-mediated effects on ER transcriptional output, thereby showcasing the power of analyzing OPS data with SCALLOPS.

## Discussion

OPS methods have enabled the linkage of genetic perturbations to rich, high-dimensional optical cellular phenotypes at a massive scale. However, the terabyte-scale image volumes produced by these methods have created significant data analysis bottlenecks, primarily in image stitching, cross-magnification registration, and scalable feature extraction, which have limited their practical application. We have developed SCALLOPS as a unified, high-performance, and modular computational framework to address these specific challenges with a robust, cloud-native solution.

A key differentiator of SCALLOPS is its integrated, "well-centric" stitching and registration architecture, which fundamentally addresses the limitations of previous tile-based workflows, such as OPS package^16^ and the recently released Brieflow^20^. SCALLOPS first stitches all tiles into a seamless well-level mosaic, thereby avoiding the problem of double-counting cells at tile overlaps. SCALLOPS then performs an automatic well-level registration across multi-magnification images, avoiding the tedious, expert-based manual process of selecting anchor points for triangle hashing registration^16,20^. PERISCOPE^13^ and recently released STARcall^31^ also adopt a stitching and registration framework. However, the stitching and registration algorithms they used are less optimized than SCALLOPS’. PERISCOPE uses FijiIS^21^ for stitching, which was shown to have worse performance than MIST^22^, while the SCALLOPS stitcher performed comparably to or better than MIST (**Extended Data Fig. 1d**). STARcall also developed a new stitcher, but it only considers phase correlation for aligning tile overlaps, uses an unweighted objective function for global optimization, and does not provide radial correction, which was shown to improve stitching quality (**Extended Data Fig. 1e**). For registration, both PERISCOPE and STARcall adopt a translation only strategy, while SCALLOPS uses a landmark-based initialization followed by a fine-grained alignment with the robust ITK elastix library^29^. The translation only strategy yielded worse registration quality with respect to bounding box correlation compared to the SCALLOPS’ strategy (**Extended Data Fig. 2d**). In addition, SCALLOPS features a computationally efficient implementation for accelerating cell morphology feature extraction and a cloud-native, WDL-based framework for horizontal scalability. Another tool, CellPaint-POSH^11^ is distinct in that the experimental design and computational workflow are coupled. It reconstructs a well-level coordinate system from tiles solely based on stage positions and registers across phenotypic and ISS modalities based on image crops at the center of wells and a phase correlation algorithm. Its registration framework is unlikely to work well on the data presented in our study without an extraordinarily accurate imaging/experimental setup. However, CellPaint-POSH features deep-learning algorithms for base calling and feature extraction, which SCALLOPS currently does not provide. We summarized the differences between SCALLOPS and alternative tools in **Supplementary Table 4**.

The methodological innovations of SCALLOPS translated novel experimental capabilities directly to higher-quality biological data. When re-analyzing the public PERISCOPE dataset, SCALLOPS both processed the data more efficiently and yielded superior results. In processing PerturbView data, SCALLOPS’ precise registration and spot detection in dense nuclear regions allowed us to accurately quantify p65 translocation, revealing a novel regulatory role for *Prkar1a* under TNF-*α* stimulation. Our unique frameshift reporter screen provided a key validation by a strong proxy for ground-truth, confirming the high reliability of SCALLOPS’ guide assignment and validating the potential of 4x magnification PerturbView for faster OPS data generation. SCALLOPS robustly stitched and registered these challenging 4x images, a task where other state-of-the-art stitchers such as ASHLAR fail (**Methods**). This unlocks a significant opportunity to accelerate OPS ISS data acquisition by up to around six-fold. Furthermore, our analysis established the 50% rule as an optimal filtering strategy, consistently providing the best balance between cell yield and guide assignment reliability.

We further demonstrated SCALLOPS’ power as a discovery engine by applying it to a multimodal screen investigating the mechanism of action of the ER-degrader vepdegestrant. We successfully validated the canonical PROTAC mechanism^41^, and this multimodal dataset, unified by SCALLOPS, allowed us to simultaneously uncover orthogonal biology, such as the role of the chromatin remodeler *ARID1A* in ER functional regulation, and generate novel, treatment-specific hypotheses for resistance, implicating genes such as *STK11, NEDD8,* and *KEAP1* in ER regulation. The ability to extract, align, and integrate these diverse data modalities, from protein levels (IF) and mRNA FISH counts to a comprehensive suite of engineered morphological features, within a single, scalable environment maximizes the information yield of OPS, facilitating its broader adoption as a comprehensive framework for systems biology.

Several avenues remain to further extend SCALLOPS’ capabilities and scalability. Transitioning from traditional computer vision algorithms to unified deep learning architectures that jointly perform spot detection and barcode calling could significantly enhance decoding accuracy, particularly in datasets with low signal-to-noise ratios. Additionally, modelling the effect of secondary barcodes in cells with more than one unique barcode can help to deconvolute the primary versus secondary effects from multiple gene knockouts. Furthermore, extending the framework to support FISH-based sgRNA detection methods, such as CRISPRmap^15^, as well as FISH-based gene-barcoding methods such as MERFISH^49^, will help integrate OPS with a broader range of spatial transcriptomics techniques in complex tissues. Finally, augmenting the current engineered feature set with deep learning-inferred phenotypic representations^11^ could help uncover latent morphological signatures, particularly in challenging cellular models such as neurons.

In summary, SCALLOPS provides a robust, scalable, and integrated software foundation necessary for the next-generation OPS. By solving the core computational bottlenecks, it lowers the barrier to entry for performing terabyte-scale experiments, improves data quality and reliability, and enables the deep, multimodal biological insights required to unravel complex disease mechanisms from pooled perturbation screens.

## Methods

### The SCALLOPS computational workflow and validation

The following computational workflow and parameters were used for all analyses described in this study, unless specified otherwise in the subsections below.

#### Illumination correction

Raw image tiles were first corrected for non-uniform illumination using the algorithm proposed in Singh et al.^50^. In particular, a flat-field profile (FFP) was generated by aggregating all images from a given channel (default aggregate method: mean). The profile was smoothed (default smooth: None; corresponding to smooth size is 1/20th of the image area) using a median filter and optionally rescaled using the 2nd percentile as a robust minimum (default rescale: True). While users can optionally provide a dark-field profile (DFP), the default DFP value was 0. The computed FFP and optional DFP are then applied to each tile I_raw_ using the formula: I_corrected_ = (I_raw_-DFP) / FFP.

#### Stitching

##### Stitching task

Given a set of tiles that are from one large image, the task of stitching is to find the (*y*, *x*) coordinates for each tile in a shared coordinate system so that the original image is reconstructed. The coordinates of tiles define the overlap regions between adjacent tiles, and the overlap regions of adjacent tiles are expected to be as similar as possible.

##### Metrics to define the similarity between overlapping regions

Assuming *f*(*y*, *x*) and *g*(*y*, *x*) are the mapping pixels of overlapping regions of two adjacent tiles, there are two ways of assessing the similarity: normalized cross correlation (NCC) and zero-normalized cross correlation (ZNCC). NCC is defined as

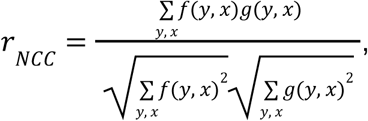

and maximizing *r*_*NCC*_ is equivalent to minimizing the normalized root-mean-square error (RMSE)^51^ of the mapped image regions in the overlapping region, which is defined as:

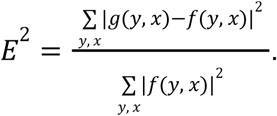

NCC can be sensitive to the mean intensity differences between the two crops. To mitigate this, the zero-normalized cross correlation (ZNCC) can be used, which is equivalent to Pearson’s correlation coefficient:

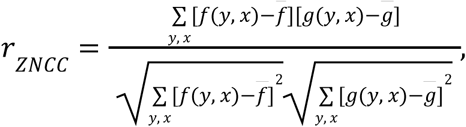

where *f̄* and *ḡ* are the mean intensity values for each mapped image region, respectively.

##### Determining the best shift for translation registration of two adjacent tiles

In typical OPS experiments, the overlap between adjacent tiles is only a small fraction of the tile (e.g. 10%). For optimal accuracy, it is therefore important to restrict the search for shift vectors around expected overlap regions instead of the whole tiles. In SCALLOPS, anticipated overlap regions from adjacent tiles are first cropped based on microscopy-provided stage positions and the image crops are then registered (see below). If the registration yields a positive ZNCC score and the updated overlap region is at least 1% (user tunable) of the tile area, an additional round of translation registration is performed based on the updated positions, aiming at further improving the registration.

##### Determining the best shift for translation registration of two image crops

To register two image crops, two Fourier transform-based algorithms are applied–one using cross-correlation^25^ and the other using phase correlation^26^–and the optimal shift vector is selected based on ZNCC scores. To enable high accurate shift vector searching and high computational efficiency, a new `find_shift` function was developed in SCALLOPS based on scikit-image’s^52^ `phase_cross_correlation` function source code. In particular, three improvements were made. First, discrete Fourier transforms are computed only once and used for both cross-correlation and phase correlation algorithms. Second, because Fourier transform-based algorithms assume the images are periodic, the shift vectors returned are not uniquely determined. Disambiguating each vector requires considering 4 symmetric directions (± △*y*, ± △*x*). In total, 8 possible shift vectors from the two algorithms are considered and the one that makes sure the overlap region is at least 1% (user tunable) of the source image crop and has the highest ZNCC score in the overlap region is selected. This improvement yields more accurate shift vectors, achieving higher ZNCC scores than scikit-image’s implementation on our data. Third, upsampling is not performed by default because upsampling did not empirically improve stitching accuracy based on our data. If users choose upsampling, it is only computed once after determining the best shift vector without upsampling, which improves computational efficiency.

##### Automatic layout determination

SCALLOPS assumes an image coordinate system where the origin is at the top left corner. However, microscope-provided stage positions are often in a different coordinate system. Thus, a procedure was implemented to automatically determine the mapping between stage positions and SCALLOPS’ coordinate system, so that it adapts to any inherent coordinate system of the microscope.

##### Stitching algorithm

The stitching algorithm is inspired by previous works such as FijiIS^21^, MIST^22^ and ASHLAR^23^. First, all tile pairs within a well are evaluated to construct an undirected adjacency graph connecting pairs whose stage-position–guided overlap relative to tile area exceeds a user-defined threshold, with edge weights given by the ZNCC score. By default, the threshold is determined as one third of the anticipated overlap region fraction over a tile, which is automatically estimated based on stage positions. A null distribution is then computed by randomly sampling 1,000 (user tunable) pairs of non-overlapping tiles and their ‘null’ shift vectors estimated from stage positions of overlapping tiles, and then computing the best ZNCC score for each sampled pair. The null distribution of ZNCC scores is approximately normal. The mean and standard deviation of this distribution are estimated robustly based on the median and median absolute deviation of the null distribution. Next, edges are filtered if they have a p-value ≥ 0.001 (user tunable) based on the null distribution or a shift in either y or x direction larger than user-defined max shift in micron (50 microns by default). Then, all connected components are detected in the resulting graph and the best positions to overlay tiles are found within each component separately (below). Lastly, the positions of each component are projected to a common coordinate system by first training a linear regressor using the tiles in the largest component as data to predict each tile’s final position from its stage position, and then applying the regressor to all tiles from other components.

To find the best positions within a connected component, the graph center is first found, defined as the vertex with minimum eccentricity. Starting from the center, the shortest paths to each vertex in the component are computed, with each edge weighted by − log *r*_*ZNCC*_. These shortest paths together define a spanning tree rooted at the graph center. The spanning tree can be viewed as a local optimal solution for the stitching problem because it does not consider all edges in the graph as constraints, and serves as an initialization for the global optimization. The global optimization procedure aims to minimize the following objective function:

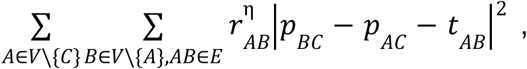

where *V* and *E* are the set of vertices and edges in the component, *C* is the graph center, *A* and *B* represent two random vertices from *V*, *r*_*AB*_ is the ZNCC score between *A* and *B*, *p*_*BC*_ and *p*_*AC*_ are position vectors relative to the center, *t*_*AB*_ is the previously estimated translation vector, and λ is a user-tunable parameter with default η = 10. 0. Compared to FijiIS, our innovation is the introduction of the *r*^η^_*AB*_ term in the objective function, which increases the weight for tile pairs with more confident overlap, indicated by high ZNCC scores, while down-weighting tile pairs with less confident overlap (e.g., tiles sharing few cells in the overlap regions and thus having small ZNCC scores). The *p* vectors are initialized using the spanning tree, which helps the algorithm converge faster, and the optimization is run for at most 5 iterations (user tunable parameter) using the `scipy.optimize.minimize` function.

##### Automatic radial distortion estimation and correction

SCALLOPS can automatically estimate radial distortion parameters from OPS images by randomly sampling 21 overlapping pairs (user tunable) from the unfiltered adjacent graph, and in parallel finding the best distortion parameter λ using the division model of distortion^24^ that maximizes the ZNCC score of the overlapping region for each pair, and finally reporting the radial distortion parameter for the whole well as the median of the 21 pairs. It then applies radial correction using the division model with the well-level parameter to all tiles and these corrected tiles are used for null distribution construction and stitching.

##### Stitching quality control (QC) report

A four-page QC report is generated per well after optimal stitching positions are determined. The first page reports the overall ZNCC score of stitched positions, defined by concatenating and squeezing all overlap regions determined by stitched positions as two long vectors and then computing the ZNCC score between the vectors. This overall ZNCC score indicates the quality of stitching and can be used to assess the performance of different stitching algorithms or optimize the stitching parameters for best performance. The first page additionally reports the radial distortion parameter and records the SCALLOPS version and stitching command used. The second page visualizes the ZNCC scores of all overlapping regions in the well. It is useful for identifying tiles without good overlaps with neighbors, which can be potentially low quality or problematic tiles. The third page visualizes the shift pattern for each tile from the staged position to the final stitched position. The last page shows a scatter plot of max shifts against ZNCC scores for all edges in the adjacent graph. Max shift for an edge is defined as the max of the absolute shifts of *y* and *x* axes estimated by SCALLOPS’ translation registration algorithm for the pair of tiles.

##### Fusing tiles into a well-level image

Tiles are fused into a seamless mosaic. To minimize edge artifacts and optical aberrations, tiles are cropped before fusion by a margin width that is dynamically estimated from the median overlap fraction of the grid. Users also have the option of setting the margin width manually. The default fusing method (blend=None) fills in the overlapping regions using pixels from the tile closest to the well center. An alternative method, setting blend to "linear", is also available, which fills in the overlapping regions using weighted average of pixel intensities from all overlapping tiles with weights proportional to each pixel’s distance from its tile edge. To handle terabyte-scale datasets efficiently, the writing process is parallelized using Dask, with memory consumption optimized by processing image channels in batches dynamically sized according to available system RAM. Tiles are written in parallel in a lock-free manner when no blending is used. When blending is selected, SCALLOPS uses shared locks, coordinated via a spatial partitioning tree, to ensure that a thread has exclusive read/write access to the region of the mosaic array it is currently processing. In addition, a mask recording the stitch boundaries is written to disk and can be optionally used later to remove cells intersecting a stitch boundary.

#### Image registration

To align stitched images (usually DAPI channel) from different cycles or modalities, all images are registered to a single ‘fixed’ reference, which is typically the first ISS cycle. All other images (e.g., subsequent SBS cycles, phenotype channels) are considered ‘moving’ images and are aligned to this reference. By default, this process computes a transformation and applies it to both the segmentation mask and the first channel (e.g. DAPI) of the moving images. The transformation is saved and applied to any other image channel and/or segmentation mask, enabling precise label transfer. By default, registration is performed on the maximum intensity projection of 3D image stacks (default z-index: "max").

The registration process begins with a landmark-based initialization (default: on) to account for large initial misalignments. In this step, template matching is performed at regular grid points (default landmark-step-size: 1,000 microns) using a defined list of template sizes (default landmark-template-padding: [750, 1,000, 1,250, 2,250] in microns). These points are filtered by a RANSAC regressor to compute a robust initial transformation. Following this initialization, the final transformation is computed using the ITK elastix library via the --itk-parameters argument. This argument accepts a list of one or more parameter sets. These can be paths to custom user-defined ITK parameter files or predefined named parameter sets. These named sets include SCALLOPS-specific sets ("rigid", "affine", "nl-100") and additional sets adapted from WSIREG^30^ (e.g., "rigid-wsireg", "affine-wsireg", "nl-wsireg", "nl-reduced-wsireg", "nl-mid-wsireg"). These sets also support metric-based suffixes such as -ams (AdvancedMeanSquares) or -anc (AdvancedNormalizedCorrelation). The default setting is a two-step registration: ["affine", "nl-100"]. This first applies an "affine" transformation (a global linear alignment for translation, rotation, shearing, and scaling). It is followed by the non-linear "nl-100" transformation on a 100-pixel grid. The final computed transformation is saved and can be applied later as needed.

#### Image segmentation

Cell segmentation is a two-step process. First, nuclei are segmented, and then cell boundaries are defined using those nuclei as seeds. By default, nucleus segmentation is performed using Stardist^53^ (--nuclei-method="stardist") with the pre-trained "2D_versatile_fluo" model. This is applied to the nucleus channel (default --nuclei-channel: 0; DAPI in our screens). Prior to segmentation, the input image is normalized using percentiles (default pmin: 3.0, pmax: 99.8). Second, cells are segmented using the propagation method^54^ (--cell-method="propagation"). This method uses a designated cell channel to create a binary cell mask, which is thresholded by default using Li’s method^55^ (alternatively “Otsu”, “Local”, or manually determined thresholds are available). The segmentation is then propagated outwards from the nucleus seeds within this cell mask. While available, background subtraction (--rolling-ball) and Gaussian smoothing (--sigma) are not enabled by default. Alternative segmentation methods such as Watershed^56,57^ and Cellpose^58^, are also supported. For processing large images, the cell segmentation is run in chunks with a default overlap of 30 pixels (--chunk-overlap). Finally, segmented objects can be filtered by size (defaults min-area and max-area are None) and a utility is provided to remove cells that intersect tile-stitching boundaries to avoid artifacts.

#### Base calling for ISS data

Base calling is a three-step process. First, candidate spots are detected using the spot detection algorithm proposed by Feldman et al.^16^. In particular, the spot detection pipeline (spot_detection_method="log") first transforms aligned ISS images using a Laplacian-of-Gaussian (LoG) filter (default σ = 1), and a maximum filter (default width=3) is applied to dilate spots. Candidate spot locations are then identified by finding local maxima (default 5x5 pixel neighborhood) in an intermediate image generated by calculating the standard deviation (std) across cycles, followed by the mean across channels. Second, crosstalk correction between channels was performed. To estimate the crosstalk matrix, all candidate spots are assembled into candidate barcodes by extracting base intensities across cycles and calculating a per-base Phred-scaled quality score as − 10*log*_10_ *P* where *P* is one minus the softmax probability of the brightest base in each cycle. A cutoff on the mean std (default threshold_peaks_crosstalk="auto") is automatically determined by sampling 500,000 candidate barcodes, defining real positives and predicted positives as candidate barcodes with a mean quality score ≥ 60 and with a mean std ≥ the cutoff respectively, and finding the optimal cutoff value that maximizes the F_0.5_ score (prioritizing precision):

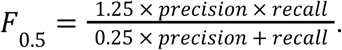

Precision and recall are defined as the ratios of true positives (instances that are both real positives and predicted positives) to the total number of predicted positives and the total number of real positives, respectively. Candidate barcodes passing this stringent mean std cutoff are used to compute a channel crosstalk matrix by pooling spots from all cycles (default crosstalk_correction_method="median")^16^. Lastly, the crosstalk matrix is applied via linear transformation to correct intensities for all candidate spots and candidate barcodes were assembled similarly as the previous step. A filtering threshold on mean std is automatically determined (default threshold_peaks="auto") by identifying the cutoff value that maximizes the F_1_ score, thereby balancing precision and recall:

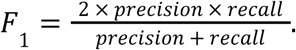

Precision and recall are defined similarly as the previous step. Candidate barcodes that pass this filter and match the whitelist (list of barcodes available in the perturbation screen) are assigned to cell labels for downstream analysis.

#### Spot detection for FISH data

Fluorescence in situ hybridization (FISH) spots were detected and quantified using the U-FISH^34^ Python library (version 1.0) with the pre-trained v1.0-alldata-ufish_c32.pth model weights. The spot detection workflow processed a multi-channel intensity image and a corresponding integer-labeled cell segmentation mask. For each fluorescence channel specified for analysis, the 2D image data were processed independently. The ufish library was used to perform image enhancement and spot calling with the following key parameters: intensity_threshold=0.06, connectivity=2, and laplace_process=True. This process yielded a list of (y, x) coordinates for all detected spots in the channel. To assign spots to cells, the coordinates of each detected spot were mapped to the cell segmentation mask. Spots with coordinates falling within a region labeled with a non-zero integer (i.e., a cell) were assigned to that cell’s label. Spots falling on the background (label 0) were discarded. Finally, the total number of spots assigned to each unique cell label was counted, generating a matrix of spot counts per cell for each channel.

#### Phenotype feature extraction

Phenotype feature extraction is performed by a custom, two-pass Dask engine designed for out-of-core computation on large-scale images. The first pass globally analyzes the label image to identify and aggregate the bounding box coordinates and centroids for all unique objects. The second pass traverses the image in parallel blocks, using the pre-computed object table to load data only for intersecting objects, calculate their features, and join the results. This engine supports a comprehensive suite of features, including standard CellProfiler-named metrics (e.g., Intensity, Texture_Haralick, Neighbors), with the specific features extracted for this work detailed in their respective sections. The engine also integrates several custom measurements, including Parameter-Free Threshold Adjacency Statistics (PFTAS), which generates 54 texture features by wrapping `mahotas.features.pftas` and applying an Otsu threshold^59^ to the object’s intensity image. For quality control, two custom metrics were implemented: ‘Bounding Box Pearson Correlation’ (correlation-pearson-box), which measures the correlation between channels using an object’s full bounding box rather than just the masked pixels, and ‘Stitching Boundary Intersection’ (intersects-boundary), which flags objects that overlap with tile-stitching boundary.

#### Workflows

Modular, scalable stitching and OPS analysis workflows were implemented in the Workflow Description Language (WDL), facilitating portable deployment across local clusters and cloud environments. The stitching workflow leverages a scatter-gather architecture to execute illumination correction and image stitching in parallel for each well and time point. The OPS workflow similarly parallelizes processing by well; furthermore, within each well, it orchestrates a dependency graph that executes independent tasks, such as segmentation, registration, and spot decoding, concurrently. To optimize performance and cost, individual tasks are containerized and configured with specific resource requests (CPU, memory) and support for preemptible instances. Cloud computing costs were estimated using AWS HealthOmics.

#### Stitching ground truth analysis

Ground truth images and evaluation code were obtained from the MIST**^22^** authors. MIST, ASHLAR, and SCALLOPS (without radial correction) were evaluated using a subset of the full dataset consisting of three replicates with 10% overlap between neighboring tiles. To quantify stitching accuracy, the primary metrics defined in the MIST benchmarking framework were employed: false positive & false negative (FP+FN), distance error (Derr) and size error (Serr), as previously detailed**^22^**. Briefly, FP+FN measures the total number of reference features (e.g., a cell colony) that are either falsely added or missed. Derr measures the Euclidean distance between the centroids of a reference feature in the stitched output and its corresponding position in the ground truth image. This metric serves as a proxy for global positioning accuracy across the well. Serr calculates the absolute percentage difference of the area of a reference feature between the stitched and ground truth images. This metric assesses local alignment quality.

### Quality control

Following feature extraction, a multi-stage filtering process was applied to the single-cell segmented data to ensure high data quality and confident barcode-phenotype linkage. This process consists of quality control filters followed by barcode purity filtering.

#### Single-cell level filters

A set of single-cell level filters was first applied to remove low-quality cells and technical artifacts. The primary filter in this set assessed the registration quality between the DAPI channels of phenotype and ISS images for each cell. This was achieved by calculating the Pearson correlation coefficient between the two imaging modalities within the entire bounding box of the cell (the correlation-pearson-box feature). Cells with a correlation score not greater than a high-confidence threshold (e.g., ≤0.9) were removed from downstream analysis. Additionally, a stitching boundary filter was applied, removing any cell mask that traversed the stitching seam in the reference phenotype.

#### Barcode purity filtering

Cells that passed single-cell level filters were then subjected to one of three barcode assignment rules of increasing stringency, all restricted to barcodes matching the whitelist: (1) Max count rule where the barcode with the highest read count is assigned to the cell, regardless of the distribution of counts for other barcodes; (2) the 50% rule, where the count of the most abundant barcode is assigned only if it is >50% of the total barcode counts for that cell; and (3) the 100% rule, where a cell is assigned a barcode only if all barcode reads detected within a cell are identical. Cells without assigned barcodes were additionally filtered out.

### Screen data retrieval, acquisition, and analyses

#### PERISCOPE A549 screen

Raw image data from all 9 plates of a genome-wide A549 Optical Pooled Screen, generated by Ramezani et al.^13^ and publicly available at the Cell Painting Gallery^60^ were reanalyzed. This dataset comprises multi-channel fluorescence images characterizing the morphological profiles of A549 cells following CRISPR-Cas9 perturbation targeting over 20,000 genes. Using the SCALLOPS pipeline with default parameters, individual tiles were stitched into a whole-well representation, phenotype and ISS images were registered, and a single-cell feature table was generated. Next, cells with low registration quality or touching stitching boundaries were filtered using the previously described single-cell level filters. Cells that passed the filters were then subjected to barcode purity filtering, retaining only those that met one of the three filtering rules above. For the correlation analysis with NGS counts, the original analysis excluded a single outlier guide, but we used all guides. For feature tables from the PERISCOPE paper and under three barcode purity filtering rules, each feature was standardized independently, dimensionality reduction was performed using PCA with 100 components, each dimension of the PCA embedding was normalized using the mean and standard deviation computed from NTCs, and guide embedding vectors were aggregated into a gene embedding by averaging. Finally, pairwise cosine similarities were computed for all gene pairs in the embedding and the recall at 5% was calculated as the fraction of top 5% gene pairs (ranked by cosine similarities) that are also in the CORUM database^35^ among all gene knockout pairs in the database.

#### PerturbView NF-κB screen

A public NF-κB p65 translocation PerturbView screen^10^ of primary mouse BMDMs was reanalyzed. In this screen, cells were transduced with a 163-gene CRISPR library, stimulated by each of three stimuli (LPS, TNF-*α* or IL-1*β*), and processed using the PerturbView protocol to image p65 (NF-κB) localization and sequence sgRNA barcodes. The three stimulations have 1, 2, and 3 wells of data respectively.

The SCALLOPS stitching workflow with default parameters was used to stitch individual tiles into a whole-well image representation. The standard SCALLOPS OPS workflow was then used to register the phenotype and ISS images and produce a single-cell feature table. A multi-step filtering process, as detailed in the "Quality control" section, was applied.

For hit calling, cells that passed single-cell level filters and the 50% barcode purity filtering rule were used. Following prior work^2,4,5,9,10^, p65 median intensities were normalized using a per-well robust z-score transformation based on the population of cells carrying non-targeting control (NTC) guides:

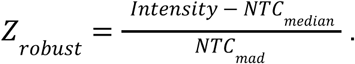

The effect size for each gene was defined as the median of guide-level median robust z-scores and p-values were computed using a bootstrap test. Specifically, for each unique cell sample size of targeting guides, a null distribution of median robust z-scores was generated by sampling NTC guides 100,000 times, resampling robust z-scores from the corresponding NTC cells with replacement to match the cell sample size, and computing the median for each iteration. A gene-level null distribution was then generated for each gene by sampling from the corresponding null distributions 100,000 times with replacement, matching the cell sample sizes of its guides, and computing the median for each iteration. The effect sizes were compared with the corresponding gene-level null distributions to determine the two-sided p-values. Benjamini-Hochberg (BH) procedure^61^ was applied to control the false discovery rate (FDR) and hits were called at FDR < 0.05. Because NTCs were used for both robust z-score normalization and null distribution generation, we considered their inclusion in the volcano plots redundant (**Fig. 2d**, **Extended Data Fig. 3d,e**). Accordingly, unlike Kudo et al.^10^, only nonessential genes are shown as controls.

#### Frameshift reporter screen

##### Benchmark dataset generation

To generate a frameshift reporter dataset for pipeline validation, an 8-cycle frameshift reporter OPS was conducted in Cas9-expressing A549 cells, following the protocol established by Kudo et al^10^. The assay used a 10-sgRNA pool (5 targeting the reporter, 5 non-targeting controls). In this system, successful perturbation triggers expression of a Hemagglutinin (HA) protein tag, which serves as a proxy for ground-truth signal to validate perturbation calls.

##### Sample processing and imaging

Ground-truth phenotype images (including DAPI and HA channels) were acquired at 20x magnification. Subsequently, ISS images were acquired from the same wells (n=3 replicates) in three distinct modalities: First, the 10x Standard was acquired using a standard OPS imaging protocol at 10x magnification. Second, the 10x PerturbView was acquired using the PerturbView protocol at 10x magnification. Finally, the 4x PerturbView was acquired using the PerturbView protocol at 4x magnification (same cells as 10x PerturbView).

##### SCALLOPS pipeline analysis and performance metrics

The standard SCALLOPS pipeline was applied to all three modalities. Due to the different ISS imaging magnifications, specific parameters were used for registration (landmark-step-size=500, landmark-template-padding 500 1000 1500 2000) and segmentation. Segmentation filters (--closing-radius 5, --nuclei-min-area 250, --nuclei-max-area 20000, --min-area 250, --max-area 20000) were then applied to remove debris and segmentation artifacts, followed by the single-cell level filters. The ‘total number of valid cells’ was defined as the cell count remaining after applying these filters. To evaluate the trade-off between cell yield and guide assignment fidelity, the barcode purity filters (max count rule, 50% rule, and 100% rule) were applied and the ‘percentage of usable cells’ was quantified as the ratio of cells passing each purity filter to the ‘total number of valid cells’.

##### Stitching 4x PerturbView tiles using ASHLAR

We tried to use ASHLAR with default parameters to stitch 4x PerturbView tiles and it failed.

#### Multimodal OPS in MCF7 cell line for vepdegestrant MoA

##### Cell culture, screening, and imaging

A doxycycline-inducible Cas9 cassette was introduced into MCF7 cells by piggybac transposon, followed by blasticidin selection. A sgRNA library was then constructed to target 146 genes involved in ER biology, along with guides targeting 12 nonessential genes and 48 NTC sgRNAs (**Supplementary Table 3**). MCF7 cells were infected with the lentiviral library at MOI of 0.1, followed by puromycin selection at 2 μg/mL for 3 days. The selected cells were then treated with doxycycline at 2 μg/mL to induce Cas9 expression. Following selection (day 6 post doxycycline induction), cells were divided into two treatment groups. The control group received DMSO, and the experimental group received 100 nM vepdegestrant. Cells were incubated for 24 hours before analysis, with each condition performed in duplicate. The subsequent experimental workflow, including cell fixation, multiplexed HCR-FISH for mRNA quantification, and immunofluorescence for ER protein, were performed as previously described^10^ with 70% ethanol permeabilization for 30 min at room temperature after fixation. Primary HCR FISH probes (Molecular Instruments) to *ESR1* (B1-488 h1/h2 amplifier), *GREB1* (B2-594 h1/h2 amplifier) and *CCND1* (B3-647 h1/h2 amplifier) were used to perform HCR FISH. For immunofluorescence, cells were incubated in primary antibody targeting ERα (Invitrogen, #MA5-14501) at 1:300 dilution for 1 h followed by donkey anti-rabbit AF488 secondary antibody (Invitrogen, #A-21206) and ConA-CF750 (Biotium, #29080) incubation at both 1:500 dilution for 45 min. After quenching the fluorescent signal with 1 mg/mL lithium borohydride (Strem, 50-901-13696) solution, ISS of sgRNA barcodes was performed for 7 cycles as previously described^2^.

##### Data preprocessing and quality control

SCALLOPS was used to process all raw images into a single-cell multimodal dataset. During cell segmentation a closing-radius parameter of 5 was used to prevent pepper noise in the mask. Nucleus size was restricted to an area between 250 and 9,600 pixels in 20x magnification, and cells between 250 and 15,000 pixels in 20x magnification, to filter potential mis-segmentations. ER protein was quantified as median nuclear fluorescence, and FISH as spot counts within whole-cell masks using SCALLOPS spot detection. Only cells that passed both the single-cell level filters and the 50% barcode purity filtering rule were retained.

##### Feature normalization, outlier removal, effect size and hit calling for the ER protein

For the median ER protein intensity feature, a local z-scoring normalization with *k* = 100 was performed to correct for fine-grained spatial variations (e.g., staining gradients), by normalizing each cell’s feature value based on its spatial neighborhood. First, a *k*-d tree was built from the centroid coordinates of all cells. For each cell, its *k* nearest neighbors (i.e., *k* = 100), including itself, were identified. The local z-score was then calculated relative to this neighborhood based on the local mean and standard deviation. Any local standard deviations of zero were set to 1 to prevent division by zero.

Next, per-perturbation outliers were filtered at the well level by removing "far out" values, defined as those falling outside the interquartile range with a *k* = 3. 0 (Tukey’s fences, “far out”).

Effect sizes were calculated as the difference between the mean of local z scores of the target gene population and the mean of the NTC population (Δµ = µ_*gene*_ − µ_*NTC*_). This metric was chosen because the mean of a non-negative random variable *X* can be expressed as the integral of its complementary cumulative distribution function (CCDF), 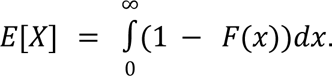 Therefore, the difference in means between two populations (e.g., gene and NTC) is mathematically equivalent to the integral of the difference between their respective CDFs:

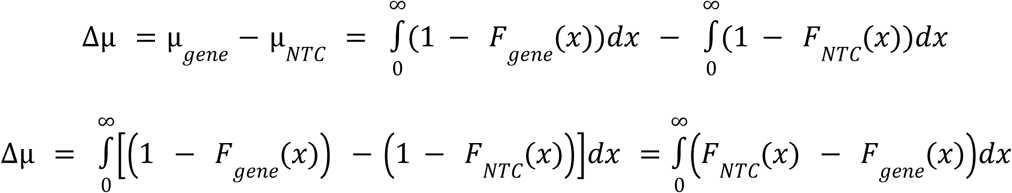

This demonstrates that Δμ represents the total signed area between the two CDF curves. This mathematical equivalence justifies its use as a robust, distribution-agnostic measure of the overall shift in the phenotype distribution. Statistical significance for this feature was assessed using a two-sided Welch’s t-test. This approach is applicable for any continuous, non-negative phenotypes (e.g., normalized protein intensity). BH procedure was applied to control FDR and hits were called at FDR≤0.05.

##### Feature normalization, outlier removal, effect size and hit calling for the FISH count data

FISH-based features (i.e., *GREB1, CCND1, ESR1*) were analyzed as raw spot counts without normalization. Next, per-feature, per-perturbation outliers were filtered at the well level using the Tukey’s fences method (*k* = 3. 0). Then a negative binomial generalized linear model (GLM) was fit for each feature separately to estimate transcript abundance, setting the NTCs as the reference baseline. The log_2_ fold change derived from the negative binomial GLM was used as the effect size, with p-values obtained directly from the model. BH procedure was applied to control FDR and hits were called at FDR≤0.05.

##### Comparative analysis of perturbation effects under vepdegestrant and DMSO

Comparative scatter plots (e.g., vepdegestrant vs. DMSO) included all genes with a significant effect (FDR≤0.05) in at least one of the conditions. Under the assumption that most genes do not have drug-specific effects, a RANSAC (RANdom SAmple Consensus) regression^28^ was used to robustly model the baseline relationship. To distinguish inliers from outliers during model fitting, the inlier threshold was set to 4.448 times the median absolute deviation (MAD) of the treatment effect sizes (∼ 3 times the standard deviation). After the final model was fitted to the identified inliers, a 99% prediction interval^62^ was computed for the regression line itself; this interval was used to highlight genes representing treatment or control-specific perturbations.

## Supporting information

Supplementary Table 1. List of features provided by SCALLOPS.

Supplementary Table 3. sgRNA design table for the ER OPS screen.

Supplementary Table 4. Comparison of SCALLOPS with related tools.

## Data availability

Raw imaging and processed data generated in this study will be available on Image Data Resource (https://idr.openmicroscopy.org/). All other data supporting the findings of this study are available from the corresponding author upon reasonable request.

## Code availability

The SCALLOPS software is available as a Python package in the Python Package Index (PyPI) and can be installed via pip install scallops. Source code is publicly available on GitHub at https://github.com/Genentech/scallops. Documentation and tutorials are publicly available at https://scallops.readthedocs.io/en/latest/. Post-SCALLOPS analysis code related to this manuscript will be publicly available at https://github.com/Genentech/scallops-manuscript.

## Acknowledgements

We thank L. Gaffney for help with figure preparation. We thank R. Thiermann for help in early benchmarking efforts of SCALLOPS on real data. We thank R. Rana for his early contribution to feature extraction code. We thank Meraj Ramezani and Sara Mostafavi for their constructive feedback on the manuscript. We thank the MIST authors for sharing their ground truth images and evaluation code, which were used for evaluating stitching algorithms. OpenAI’s ChatGPT and Google’s Gemini were used to assist with language editing and polishing of the manuscript.

## Author contributions

J.Gould, J.S.H. and B.L. conceived the study. J.Gould and J.S.H developed the SCALLOPS software. T.K. designed and performed the frameshift reporter OPS experiments, with inputs from J.Gould, J.S.H., E.L. and B.L.. P.W., C.M. and A.S. designed the vepdegestrant OPS screen experiment. P.W. and J.Guan performed the experiment. J.Gould and J.S.H. analyzed data. J.Gould, J.S.H. and B.L. interpreted data. A.Z. contributed to the SCALLOPS code base and advised on the statistics. X.G. and A.A.W. supported the analysis. J.S.H., J.Gould, and B.L. wrote the paper, with inputs from all authors. T.B., O.R.R., D.R., A.R. and B.L. supervised the work.

---

**Extended Data Figure 1.**
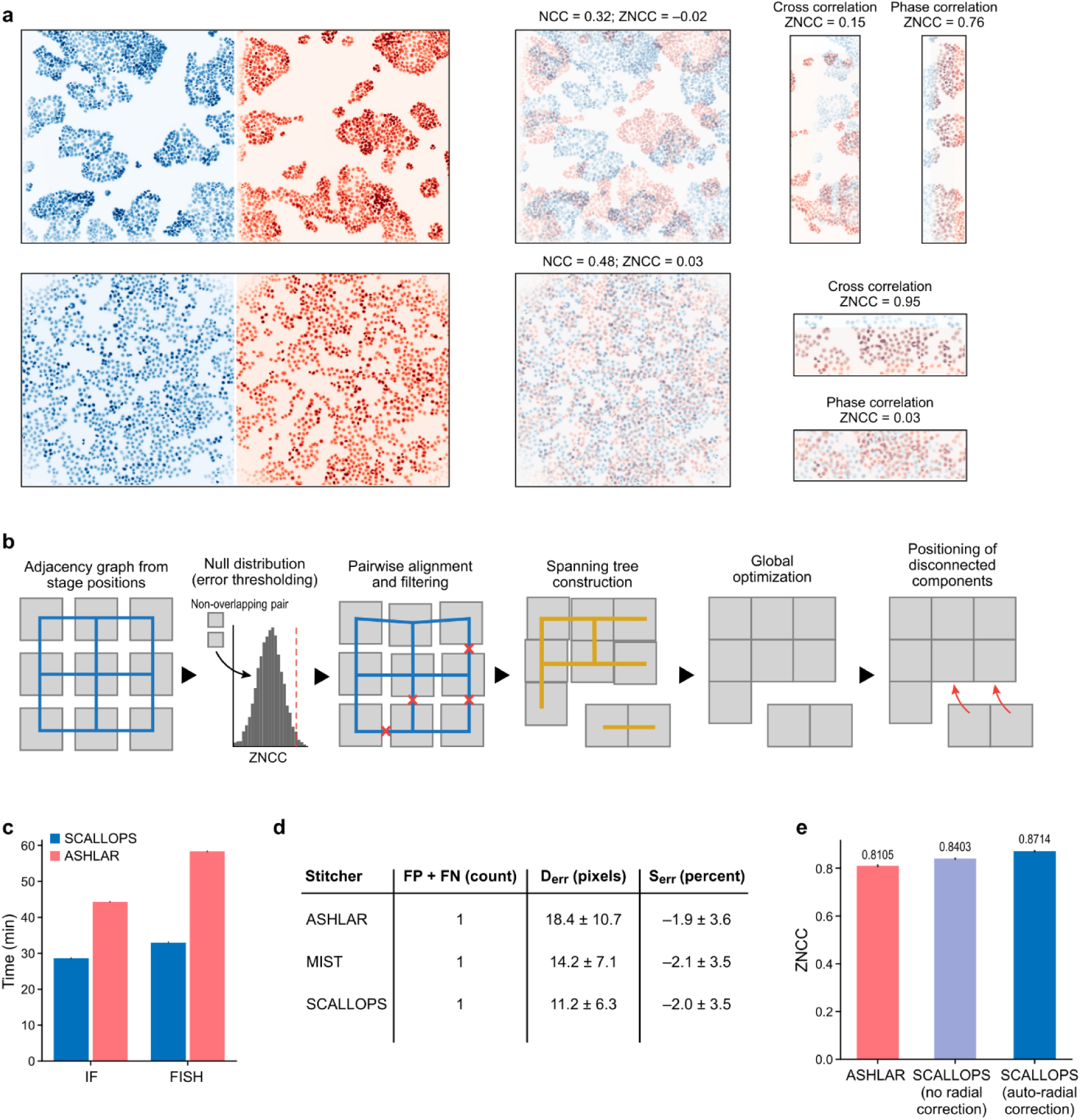
SCALLOPS stitching algorithm design and performance benchmarks. **a,** Comparison of alignment methods across overlapping regions in two distinct datasets: ER (top) and Funk et al.^9^ (bottom). For each, two contiguous tiles (left) and their full overlap (middle) are shown alongside results for cross correlation and phase correlation (right). **b,** Schematic of the SCALLOPS stitching workflow. From left: construction of a filtered adjacency graph from overlapping tiles, local and global optimization to determine relative positions for each connected component, and projection of all components into a shared coordinate system via linear regression. **c,** Stitching runtime comparison between SCALLOPS and ASHLAR. Processing times are shown for IF (3 channels) and FISH (4 channels) panels. Error bars: standard error of the mean (SEM). **d,** Performance benchmarking on the MIST ground truth dataset. SCALLOPS (without radial correction) is compared against MIST and ASHLAR across three metrics: FP + FN (left), D_err_ (middle), and S_err_ (right). Lower values indicate better performance. **e,** Quantitative assessment of stitching accuracy across four wells of ER data. Mean ZNCC scores are shown for ASHLAR, SCALLOPS without radial correction, and SCALLOPS with radial correction. Error bars: SEM.

**Extended Data Figure 2.**
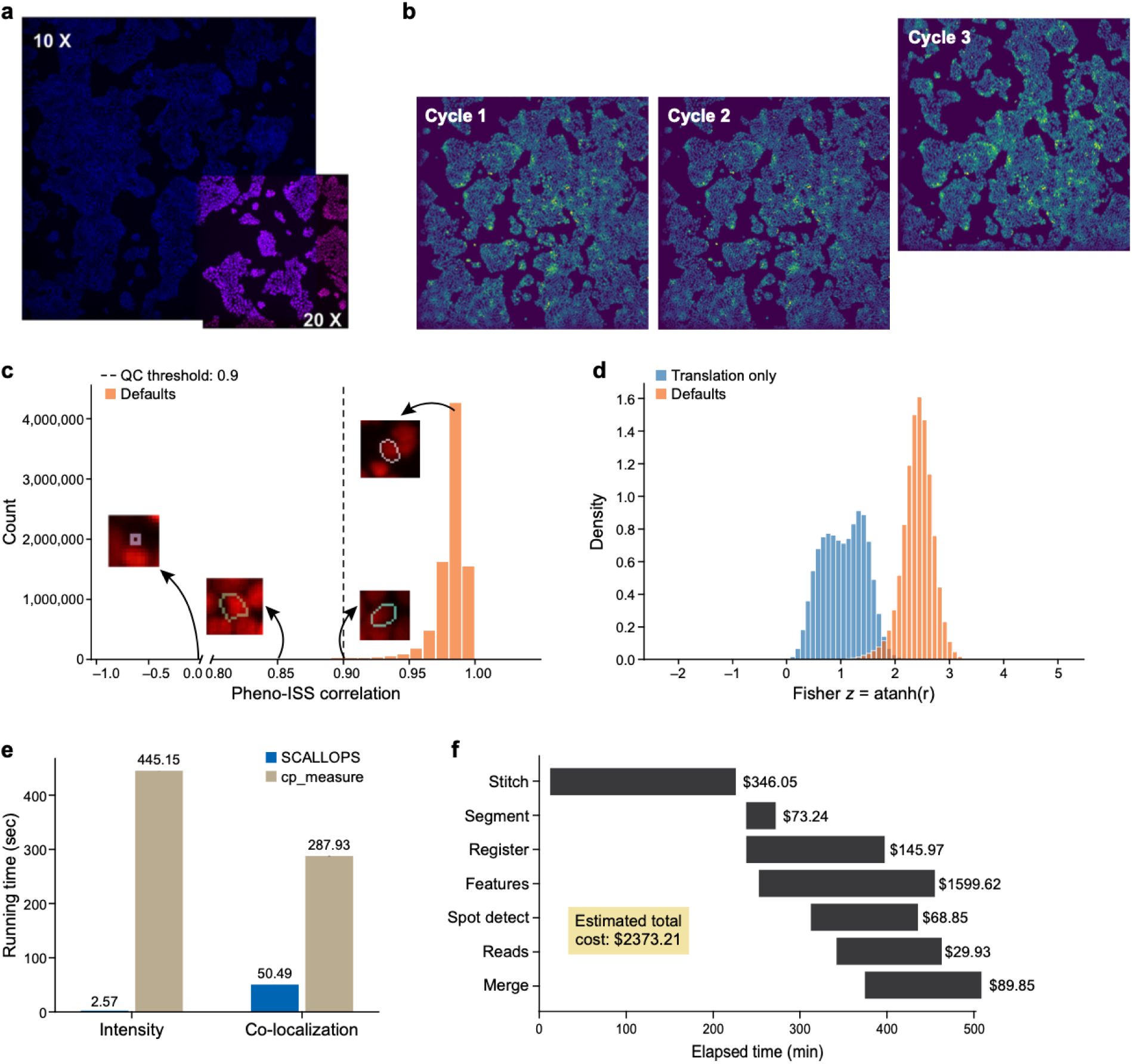
Registration diagnostics and computational scalability. **a,** Representative raw images illustrating the spatial relationship between a 20x phenotypic tile (magenta) and a 10x ISS tile (blue). **b,** Demonstration of cycle-to-cycle misalignment. Raw images of the same field of view across three consecutive ISS cycles (cycle 1-3) showing a physical shift in cycle 3. **c,** Quality control metric for registration. Distribution of bounding box Pearson correlations for DAPI intensities between registered phenotype and ISS images. Insets: representative crops showing misalignment (correlation = 0), marginal alignment (0.85), the filtering threshold (0.90), and high-fidelity alignment (0.98). **d,** Comparison of registration strategies. Probability density of Fisher-transformed bounding box Pearson correlations for cells registered using a ‘translations only’ method (blue) or the full SCALLOPS two-stage pipeline (orange). **e,** Feature extraction efficiency. Runtime comparison for intensity and colocalization features between SCALLOPS and cp_measure (calculated from 607 cells in a 1,480 x 1,480 pixel region). **f,** Cloud pipeline performance. Total runtime (x axis) and computational cost (bar labels) for each stage (y axis) of the SCALLOPS WDL pipeline on AWS for the full PERISCOPE A549 dataset (9 plates).

**Extended Data Figure 3.**
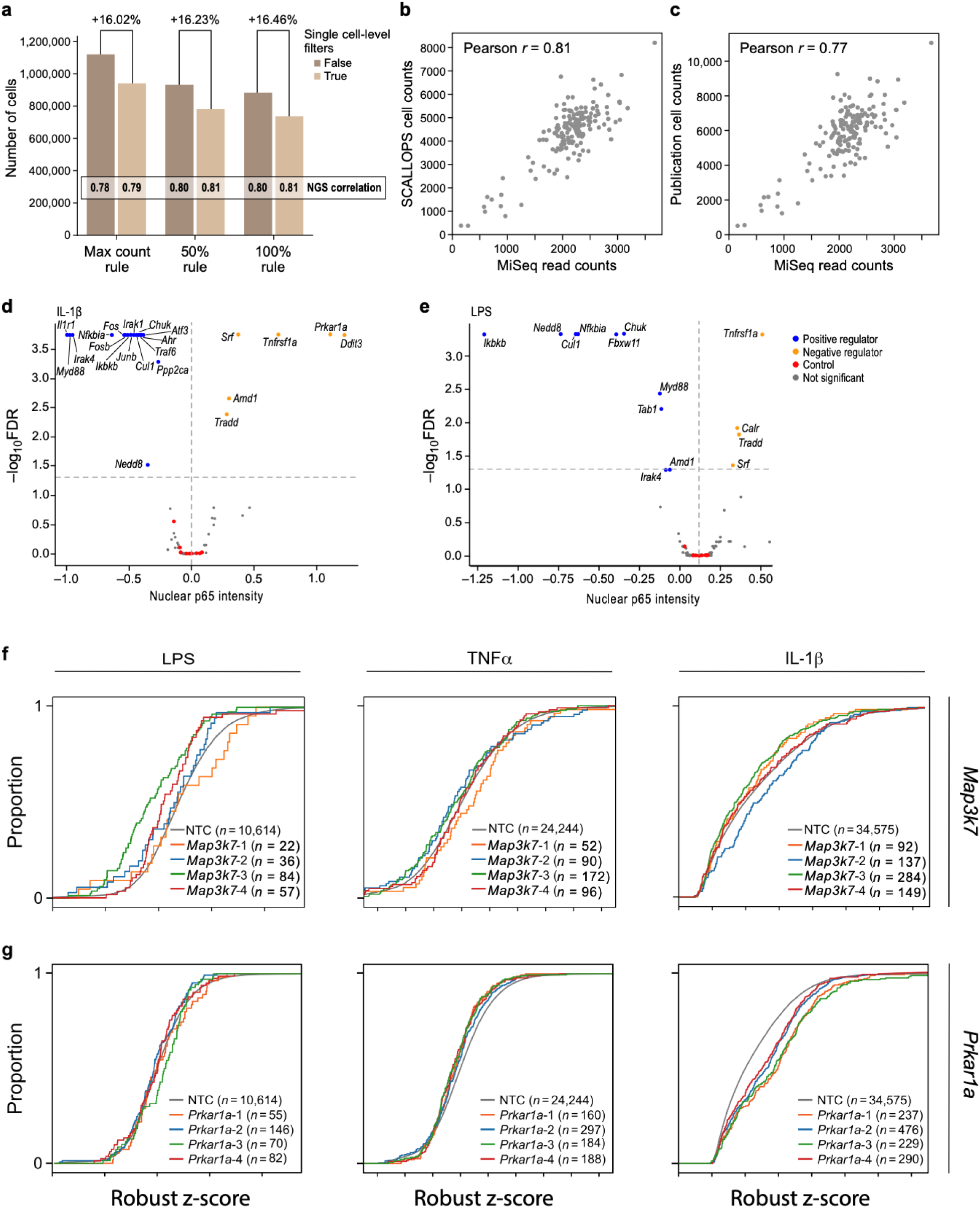
Identification of known and novel regulators in the BMDM PerturbView screen. **a,** Impact of single-cell level filters on cell recovery. Number of usable cells in the BMDM PerturbView screen using SCALLOPS with or without applying single-cell level filters across three barcode purity filtering rules (max count, 50%, and 100%). Pearson’s correlations with plasmid NGS counts are shown within each bar. **b,c,** Correlation between plasmid NGS counts and cell counts for SCALLOPS (50% rule) (**b**) and the original analysis (**c**). **d,e,** Identification of context-specific hits. Significance (-log_10_(FDR), y axis) and effect sizes (defined as the median of guide-level medians of normalized nuclear p65 intensity, x axis) for each gene (dot) under IL-1β (**d**) or LPS (**e**) stimulation, colored as positive regulators (blue; FDR<0.05), negative regulators (orange; FDR< 0.05), non-significant (grey; FDR≥0.05) or nonessential gene controls (red). **f,g,** Cumulative distribution functions (CDFs) of robust z-scores of median nuclear p65 intensity for NTCs (grey) and guides targeting *Map3k7* (**f**) and *Prkar1a* (**g**) under LPS (left), TNF-*α* (middle), and IL-1*β* (right) stimulation.

**Extended Data Fig. 4.**
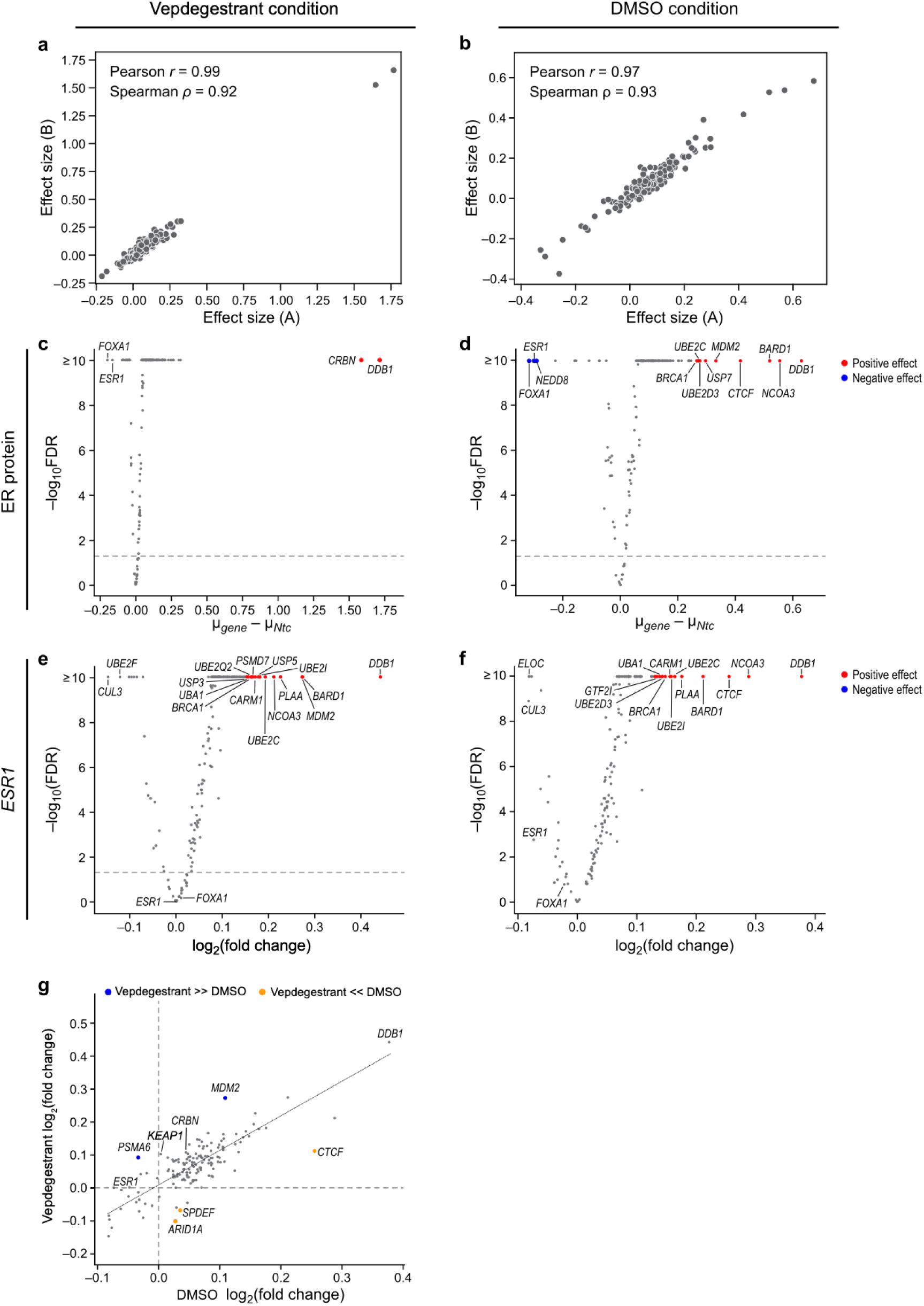
Reproducibility and hit identification in the vepdegestrant optical pooled screen. **a,b,** Reproducibility of gene-level knockout effects on ER protein abundance between plates A (x axis) and B (y axis) under vepdegestrant (**a**) or DMSO (**b**) treatment. Pearson’s and Spearman’s rank correlation coefficients are indicated in the top left of each panel. **c,d,** Gene-level knockout effects on ER protein abundance under vepdegestrant (**c**) or DMSO (**d**) treatment. Significance (-log_10_(FDR), y axis) and effect size (difference in mean normalized median nuclear ER intensity between genes and the NTC, x axis) for each gene (dot). Labeled genes: hits with FDR≤0.05 and effect sizes > 2 s.d. from the mean, or of interest (*ESR1*, *FOXA1*). **e,f,** Gene-level knockout effects on *ESR1* transcript abundance under vepdegestrant (**e**) or DMSO (**f**) treatment. Significance (-log_10_(FDR), y axis) and log_2_ fold change (x axis) for each gene (dot). Labeled genes: hits with FDR≤0.05 and log_2_ fold change > 2 s.d. from the mean or of interest (*ESR1*, *FOXA1*, *CUL3*, *UBE2F*, *ELOC*). **g**, Treatment-dependent gene-level knockout effects on *ESR1* transcript abundance. Genes (dots) with significant (FDR≤ 0.05) effects on *ESR1* transcript abundance in DMSO (x axis) or vepdegestrant (y axis) treatment. Colored points: hits with higher effect in vepdegestrant (blue) or DMSO (orange) conditions (outside the RANSAC regression 99% prediction interval, dotted black line). Genes of interest (*CRBN*, *DDB1*, *ESR1*, *KEAP1*) are marked. *FOXA1* was not significant and is not shown.

**Extended Data Fig. 5.**
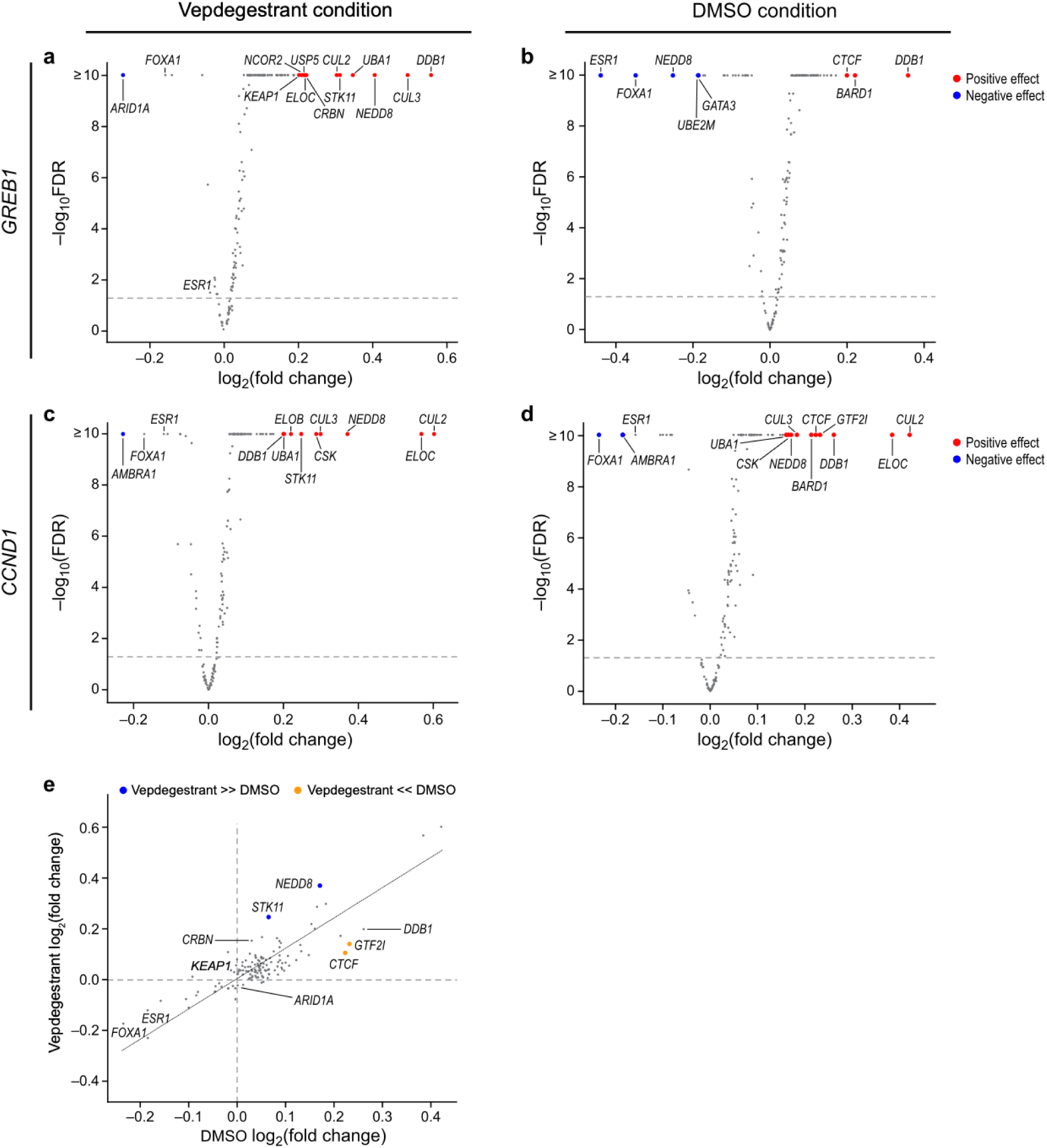
Gene-level knockout effects on *GREB1* and *CCND1* transcript abundance. **a,b,** Gene-level knockout effects on *GREB1* transcript abundance under vepdegestrant (**a**) or DMSO (**b**) treatment. Significance (-log_10_(FDR), y axis) and log_2_ fold change (x axis) for each gene (dot). Labeled genes: hits with FDR≤0.05 and log_2_ fold change > 2 s.d. from the mean or of interest (*ESR1*, *FOXA1*). **c,d,** Gene-level knockout effects on *CCND1* transcript abundance under vepdegestrant (**c**) or DMSO (**d**) treatment. Significance (-log_10_(FDR), y axis) and log_2_ fold change (x axis) for each gene (dot). Labeled genes: hits with FDR≤0.05 and log_2_ fold change > 2 s.d. from the mean or of interest (*ESR1*, *FOXA1*). **e,** Treatment-dependent gene-level knockout effects on *CCND1* transcript abundance. Genes (dots) with significant (FDR≤0.05) effects on *CCND1* transcript abundance in DMSO (x axis) or vepdegestrant (y axis) treatment. Colored points: hits with higher effect in vepdegestrant (blue) or DMSO (orange) conditions (outside the RANSAC regression 99% prediction interval, dotted black line). Genes of interest (*ARID1A*, *CRBN*, *DDB1*, *ESR1*, *FOXA1*, *KEAP1*) are marked.

